# Transitions between cognitive topographies: contributions of network structure, neuromodulation, and disease

**DOI:** 10.1101/2023.03.16.532981

**Authors:** Andrea I. Luppi, S. Parker Singleton, Justine Y. Hansen, Danilo Bzdok, Amy Kuceyeski, Richard F. Betzel, Bratislav Misic

## Abstract

Patterns of neural activity underlie human cognition. Transitions between these patterns are orchestrated by the brain’s network architecture. What are the mechanisms linking network structure to cognitively relevant activation patterns? Here we implement principles of network control to investigate how the architecture of the human connectome shapes transitions between 123 experimentally defined cognitive activation maps (cognitive topographies) from the NeuroSynth meta-analytic engine. We also systematically incorporate neurotransmitter receptor density maps (18 receptors and transporters) and disease-related cortical abnormality maps (11 neurodegenerative, psychiatric and neurodevelopmental diseases; *N* = 17 000 patients, *N* = 22 000 controls). Integrating large-scale multimodal neuroimaging data from functional MRI, diffusion tractography, cortical morphometry, and positron emission tomography, we simulate how anatomically-guided transitions between cognitive states can be reshaped by pharmacological or pathological perturbation. Our results provide a comprehensive look-up table charting how brain network organisation and chemoarchitecture interact to manifest different cognitive topographies. This computational framework establishes a principled foundation for systematically identifying novel ways to promote selective transitions between desired cognitive topographies.

## INTRODUCTION

The brain is a complex system of interconnected units that dynamically transitions through diverse activation states supporting cognitive function [18, 19, 24, 25, 129, 131]. Large-scale, noninvasive techniques like functional magnetic resonance imaging (fMRI) provides a way to map activation patterns to cognitive functions [7, 32, 76, 93, 115, 159]. Healthy brain function requires the ability to flexibly transition between different patterns of brain activation, to engage the corresponding cognitive functions in response to environmental and task demands. In turn, the neurophysiological dynamics of the human brain are both constrained and supported by the network organisation of the structural connectome: the white matter fibers that physically connect brain regions [3, 61, 62, 96, 138]. However, the exact mechanisms by which the brain’s network architecture shapes its capacity to transition between cognitively relevant activation patterns remain largely unknown, and an intense focus of inquiry in neuroscience [6, 17, 73, 130].

Network control theory is a computational paradigm that explicitly operationalises how the architecture and dynamics of a network supports transitions between activation states [52, 73, 80]. Originally developed in the physics and engineering literature [80, 81, 126], network control theory conceptualises the state of a dynamical system at a given time as a linear function of three elements: (i) the previous state, (ii) the structural network linking system units, and (iii) input injected into the system to control it.

In the context of the brain, such input can intuitively take the form of task modulation [14, 16, 51] or other perturbations from the environment, but potentially also pharmacological or direct electromagnetic stimulation [72, 91, 124, 134], or endogenous signals from elsewhere in the brain [92]. This approach is widely applicable across the breadth of neuroscience, from *C. elegans* and *drosophila* [73, 158] to rodents and primates [52, 73], and across human development [78, 106, 140], health, and disease [11, 14, 16, 58, 107, 134, 165, 166].

In humans, network control can be used to study the transition between brain states. Such state-to-state transitions can be formalised as a dynamical process that unfolds over the connectome’s network architecture, reconstructed from diffusion-weighted imaging (DWI). Of particular relevance is the quantification of control energy. Control energy refers to the magnitude of input that needs to be provided to the system in order to drive its trajectory from an initial state to a desired target state [52, 80]. In the context of transitions between brain states, the cost of transitions may correspond to the magnitude of exogenous stimulation (e.g. transcranial magnetic stimulation, deep brain stimulation, intracranial stimulation [36, 91, 97, 134] or the dose of a pharmacological intervention [16, 49, 124]), but also to endogenous effort, as reflected by cognitive demand [90, 92].

When seeking to operationalise this promising framework, conventional studies on control energy in the human brain have consistently adopted one of two strategies for defining brain states. One strategy is to define brain states as co-activations of cognitively relevant brain circuits, operationalised as the canonical intrinsic connectivity networks of the brain [14, 22, 51, 58, 70]. The downside of this approach is that intrinsic networks identified from functional MRI are limited in number (usually only 7–8 [20, 23, 110, 129, 163]), providing a correspondingly limited repertoire compared with the space of possible functional activation patterns. The second strategy typically involves defining brain states as random activation patterns [58, 73], whose number is then virtually limitless, but at the expense of being cognitively ambiguous.

Here we overcome these challenges by investigating how network architecture supports transitions between cognitive topographies. We define cognitive topographies (i.e., cognitively relevant brain states) as meta-analytic patterns of cortical activation pertaining to over 100 cognitive terms, obtained by aggregating over 15 000 functional MRI studies from the NeuroSynth atlas [159]. This approach represents a large-scale generalisation of recent work that defined cognitively relevant brain states in terms of task-based fMRI contrast maps [16, 68].

In addition to generalising the set of possible start and target states under consideration, we also provide two key extensions to the scope of the control inputs under investigation. First, we consider the potential role of deficiencies in the capacity of brain regions to act as sources of endogenous control signals, due to pathology. We operationalise this using cortical thickness abnormalities for 11 neurological, psychiatric, and neurodevelopmental disorders from the ENIGMA consortium, summarising contrasts between 17 000 patients and 22 000 controls [56, 77, 141, 142]. Second, we investigate the potential role of pharmacological control on cognitive transitions by defining inputs as the regional expression of 18 neurotransmitter receptors and transporters, quantified from *in vivo* positron emission tomography (PET) scans in > 1 200 subjects [55]. Overall, we combine multiple databases from different neuroimaging modalities (functional MRI, diffusion MRI tractography, cortical morphometry, PET) to investigate how the brain’s network architecture shapes its capacity to transition between a large number of experimentally defined cognitive topographies, and how this capacity can be reshaped by perturbing control inputs due to disease or pharmacology.

## RESULTS

Network control allows us to ask how brain network structure supports transitions between cognitively-relevant brain states [52, 92] (Fig. 1a). We define the network whose activity is to be controlled as the human structural connectome (obtained as a consensus of *N* = 100 Human Connectome Project [148] subjects’ connectomes reconstructed from diffusion MRI tractography; see *Methods*). We define the cognitive topographies as meta-analytic brain activation patters from the NeuroSynth atlas (*Methods*). With definitions of the network and its state in hand, we consider the problem of network controllability: how can the system be driven to specific target states by internal or external control inputs (Fig. 1b)? Beginning with uniform inputs applied to all regions, we are interested in the relative energetic cost of transitions between different cognitive topographies (Fig. 1c).

**Figure 1.**
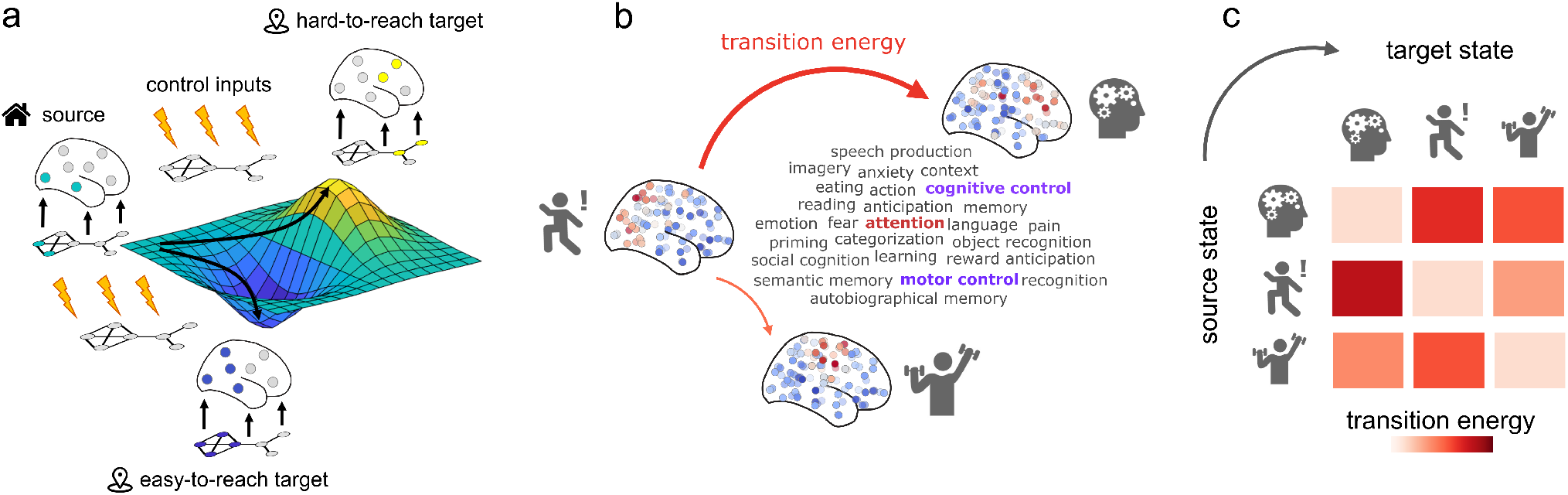
Network control with cognitive topographies. **(a)** Functional brain activity (coloured nodes are active, grey nodes are inactive) evolves through time over a fixed network structure (displayed below the brains). From a given starting configuration of activity (green), some alternative configurations are relatively easy to reach in the space of possible configurations (valley, in blue), whereas others are relatively difficult to achieve (peak, in yellow). To reach a desired target configuration of activity, input energy (represented by the lightning bolt icons) can be injected locally into the system, and this energy will spread to the rest of the system based on its network organisation. **(b)** We define states as 123 meta-analytic activation maps from the NeuroSynth database. We then use network control theory to quantify the cost of transitioning between these cognitive topographies. **(c)** Systematic quantification of transition cost between each pair of cognitive topographies results in a look-up table mapping the energy required for each transition.

### Transitions between cognitive topographies

We first evaluate the control energy required to transition between each pair of cognitive topographies (“brain states”) from NeuroSynth [159]. The NeuroSynth meta-analytic engine provides meta-analytic functional activation maps associated with 123 cognitive and behavioural terms from the Cognitive Atlas [109], ranging from general terms (“attention”, “emotion”) to specific cognitive processes (“motor control”, “autobiographical memory”), behavioural states (“eating”, “sleep”), and emotional states (“fear”, “anxiety”). Each map is a vector encoding the statistical strength of differential activation for task-based neuroimaging studies that include a cognitive term versus studies that do not include that term, at each spatial location, based on the published literature. Although NeuroSynth uses activation maps as the inputs for its term-based meta-analysis, its outputs are not activation maps per se, but rather they reflect statistical association tests.

Applying control inputs uniformly to all brain regions, we compute the optimal energy cost for each of the 15 129 possible transitions between cognitive topographies. We find that optimal control energy can vary by nearly 10-fold across different transitions (Fig. 2a–c). As a result of these differences, for several combinations of source and target cognitive topographies (36%) a direct transition is not the most energy-efficient. Rather, the control energy required to transition between them can be reduced if an intermediate transition is made to some other state (Fig. S1).

**Figure 2.**
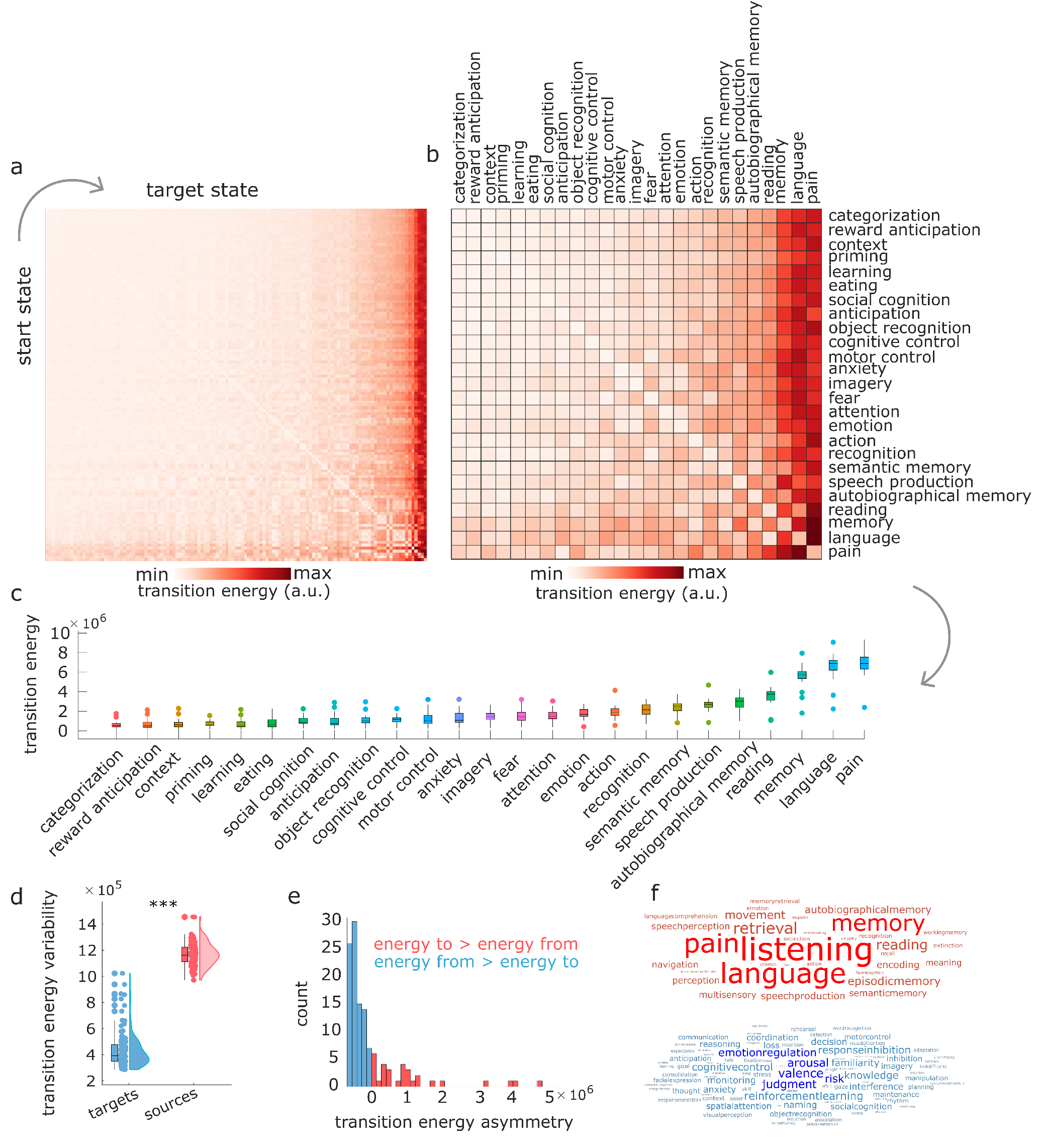
Quantifying transitions between cognitive topographies. **(a)** Transition cost (energy) between each pair of 123 cognitive topographies (“states”) from NeuroSynth. Rows indicate source states, columns indicate target states. **(b)** Same as (a), but showing only an arbitrary subset of 25 out of 123 NeuroSynth states for visualisation purposes. Matrices are sorted by increasing cost across both rows and columns. **(c)** Distributions of the cost to transition to each cognitive topography from every other cognitive topography. **(d)** Variability (standard deviation) of transition energy is greater along the column dimension (target states) than along the row dimension (source states). **(e)** Histogram of the difference in transition cost between reaching each state (averaging across all possible source states) and leaving each state (averaging across all possible target states). Positive values indicate greater cost to reach a state than to leave it, whereas negative values indicate the reverse. **(f)** Word clouds display the NeuroSynth terms that are more difficult to reach than to leave, on average (red), or more difficult to leave than to reach, on average (blue). Word size reflects ranking.

The transition energy between each pair of states correlates with the Euclidean distance between their NeuroSynth vector representations (Spearman’s *r* = 0.99, *p* < 0.001): the more distant two patterns are, the more energy will be required, and consequently the average energy to reach each target state correlates with the mean (Spearman’s *r* = 0.49, *p* < 0.001) and standard deviation (Spearman’s *r* = 0.96, *p* < 0.001) of the corresponding NeuroSynth map (Fig. S2a-d).

However, mean and variance of the NeuroSynth activation patterns are coarse descriptions of each pattern. This is because they disregard all information about the neuroanatomical distribution of activations. Additionally, Euclidean distance alone cannot fully account for the observed results: Euclidean distance is symmetric, whereas we observe that transition energy is asymmetric. Specifically, we observe that target states (columns of the matrix) exhibit greater variability than start states (rows), suggesting that the destination of a transition— the desired target topography—may play a more prominent role than the current state in determining the ease or difficulty of the transition (Fig. 2a–c). After partialling out the effect of each NeuroSynth map’s mean and standard deviation, we find that its transition cost is related to both topological features of the structural connectome (especially whether high-valued nodes are easy or difficult to reach using a diffusion process), and the map’s spatial alignment with the unimodal-transmodal cortical hierarchy (Fig. S2e-g).

We confirm the observation of transition asymmetry by showing that the variability (standard deviation) of the transition energy matrix is higher across target states (mean = 1.17 × 10^6^, SD = 8.74 × 10^4^) than across start states (mean = 4.35 × 10^5^, SD = 1.35 × 10^5^; t(244) = 50.52, *p* < 0.001, Cohen’s *d* = 6.42) (Fig. 2d). To investigate this asymmetry further, we also compute a measure of transition asymmetry between each pair of brain states *i* and *j*, as the difference in control energy required to move from *i* to *j*, versus moving from *j* to *i*. Averaging across start states provides, for each target state, a measure of whether that brain state is overall easier to reach than leave (negative values) or harder to reach than leave (positive values) from other states. As expected, this measure is positively correlated with the overall transition cost to reach a given state (Spearman’s *r* = 0.73, *p* < 0.001). We find that the majority of cognitive topographies are slightly easier to reach than to leave, but this is counterbalanced by a small number of cognitive topographies that are substantially harder to reach than to leave. In particular, hard-to-reach cognitive topographies include those pertaining to language-related cognitive operations (e.g., “language”, “reading”, “speech production”) and those pertaining to memory (e.g., “memory”, “autobiographical memory”, “semantic memory”)(Fig. 2e,f). We confirm this observation quantitatively via empirical permutation tests (1, 000 permutations): we consider the six data-driven cognitive domains identified by Beam et al. [7]: memory, cognition, inference, emotion, vision, and language. Among the terms in our list of 123 that have been assigned to one of these six domains, we find that terms pertaining to “memory” (*p* = 0.007) and “language” (*p* = 0.045) exhibit higher median value of asymmetry than would be expected by chance (“emotion”: *p* = 0.640; “inference”: *p* = 0.696; “cognition”: *p* = 0.542; “vision”: *p* = 0.730). Altogether, we find that the energetic ease or difficulty of transitioning between two given cognitive topographies appears to be primarily driven by the identity of the target state.

### Connectome wiring supports efficient cognitive transitions

Having considered how transition energy varies as a function of different origin and target cognitive topographies, we now turn our attention to the network itself. We assess how much the observed effects are due to topology and geometric embedding. To address this question, we implement two classes of null models [150].

We first consider a null model that preserves weight distribution and degree sequence [88]; and a “geometry-preserving” rewired null, preserving degree sequence and weight distribution, but also the approximate wiring cost (length of connections) [12]. For each specific null model, we re-estimate the control energy 500 times, and compare the resulting distribution of all-to-all mean transition energy against the distribution obtained using the empirical structural connectome of the human brain.

We find that the human brain outperforms both null models: transitions are significantly less energy-expensive on the human connectome than on either null (all Fig. 3), suggesting that the unique wiring architecture of the human connectome supports efficient transitions between cognitively relevant topographies. We also find a significant difference between the nulls: rewired networks are more energy-efficient when they preserve both the degree sequence and the geometric properties of the human connectome, than when only the degree sequence is preserved. In other words, the geometric embedding of the human connectome accounts for a substantial portion of its energy-efficiency—but not all of it. We also show that these observations, obtained from a consensus connectome, remain true at the single-subject level: subject-level matrices of transition energy correlate with the group-level matrix, and individual connectomes outperform corresponding degree-preserving and degree- and cost-preserving nulls (Fig. S3 and Table S1).

**Figure 3.**
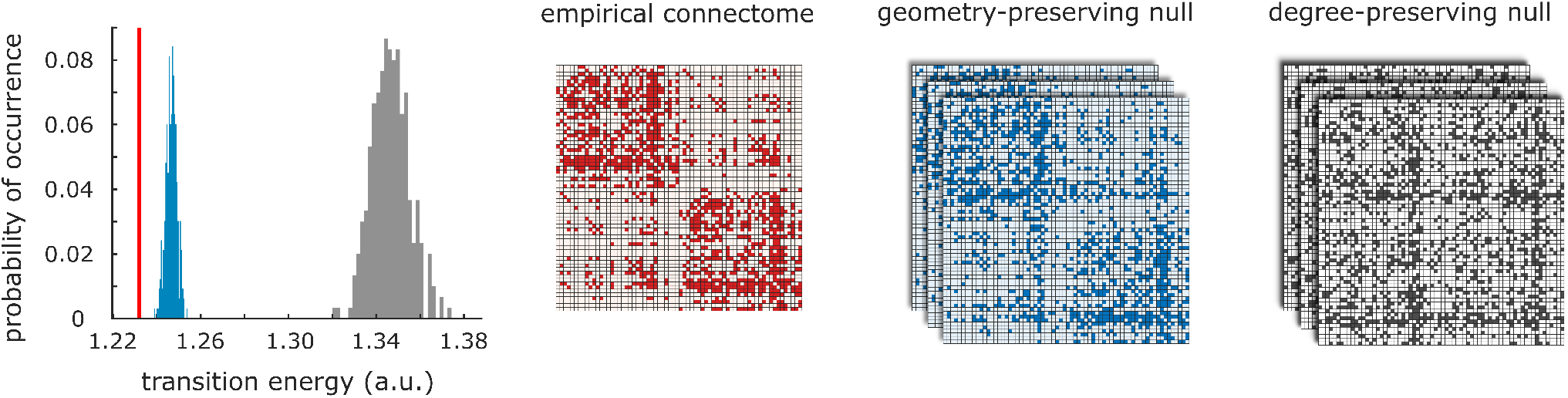
Role of network topology in supporting transitions between cognitive topographies. Degree-preserving randomised null models (grey) and null models that preserve the exact degree sequence and the approximate length distribution (blue) are significantly less favourable than the empirical human connectome (red) to support transitions between cognitive topographies. Note that all networks are weighted and have the exact same distribution of weights, but they are displayed here as binary to highlight the respective similarities and differences in topology.

### Effect of disease on transitions

Up to this point, we investigated both the role of the transition sources and targets, and the role of the underlying network. We now turn to the remaining element of the network control theory as a framework for dynamics on neuronal networks: the control inputs. To this end, we consider how pathology-associated variations in cortical thickness may impact the connectome’s ability to facilitate transitions between cognitive topographies. We focus on cortical thickness by making the simplifying assumption that the amount of input energy that a region is capable of injecting into the network is a function of its total number of excitatory neurons (since most interregional projections are known to originate from excitatory neurons)—and that this, in turn, is related to cortical thickness (since most neurons are excitatory). This approach is conceptually related to the approach recently employed by Singleton and colleagues [124], who applied non-uniform control inputs according to the normalised density of the serotonin 2A receptor expressed in each region (as quantified by PET).

We consider different patterns of abnormal cortical thickness associated with 11 neurological diseases and neuropsychiatric disorders (collectively referred to as “disorders” here, for brevity), as quantified by the ENIGMA consortium and recent related publications [56, 77, 141, 142]. For each of 11 ENIGMA disorders, we modulate the regional control input according to the regional pattern of increases or decreases in cortical thickness associated with that disorder (Fig. 4).

**Figure 4.**
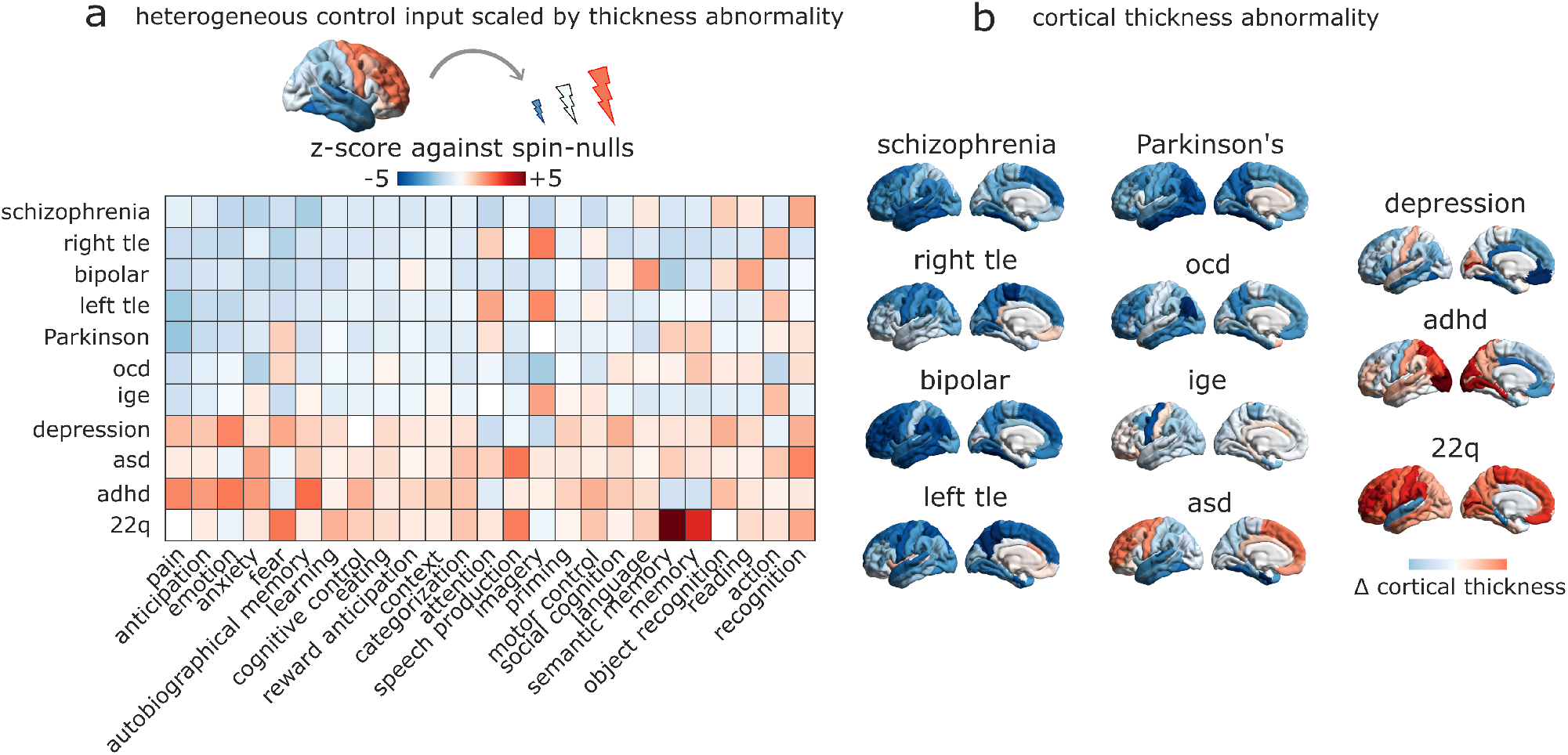
Simulating the effect of disease-associated cortical atrophy on transitions. **(a)** The effect of cortical abnormality is modelled by changing the control input provided by each region in proportion to the extent of its cortical thickness alteration from healthy controls (Cohen’s *d*): atrophied regions exert less input, and regions of increased thickness exert greater input. Heatmap shows how each disease reshapes the average transition energy required to reach a given cognitive topography from all other cognitive topographies, presented as a *z*-score against a null distribution of randomly rotated cortical thckness abnormality maps with preserved spatial autocorrelation and the same increases and decreases in cortical thickness, but occurring at different neuroanatomical locations. **(b)** The changes in cortical thickness associated with each disease are shown on the cortical surface. adhd = attention deficit/hyperactivity disorder; asd = autistic spectrum disorder; ocd = obsessive-compulsive disorder; ige = idiopathic generalised epilepsy; right tle = right temporal lobe epilepsy; left tle = left temporal lobe epilepsy.

In particular, we seek to disentangle the role of overall changes in cortical thickness, versus their specific neuroanatomical distribution. To this end, and to remove the potential confound of the mean and variance of each distribution, we compare the transition energy associated with each pattern of cortical thickness, with randomly rotated versions of the same pattern, preserving the original brain map’s spatial autocorrelation and the overall distribution of values, but randomising the neuroanatomical locations [86]. For each disorder, we *z*-score the empirical control energies against the resulting null distributions of transition energies, separately for each target cognitive topography. For a given combination of disorder and target state, a negative *z*-score indicates that the real disorder-associated pattern requires overall less energy than the null patterns, to induce a transition to the target state in question. A positive *z*-score indicates the opposite. Overall difference in the cost of all pairwise transitions, compared with baseline, is shown in (Fig. S4): as expected, since most alterations in cortical thickness are reductions, the overall control input is diminished, and thus the transition cost is higher.

Our results show that for some disorders, such as schizophrenia, bipolar disorder, or the temporal lobe epilepsies, the neuroanatomical distribution of cortical abnormalities is such that transition costs are on average lower than would be expected by only considering equivalent but randomly distributed changes in cortical thickness. In contrast, other conditions such as autism, ADHD, depression and 22.q syndrome incur transition costs that exceed what would be expected based on randomly distributed cortical abnormalities (Fig. 4).

### Effect of pharmacological manipulations on transitions

Finally, mimicking the effects of endogenous or pharmacological modulation, we modulate control inputs in proportion to the regional expression of different neurotransmitter receptors and transporters, quantified from in vivo PET. We consider a total of 18 PET maps [55]. This allows us to evaluate how each receptor’s distribution favours transitions towards different cognitive topographies. As above, we account for the overall distribution of values in each receptor map by comparing it with randomly rotated null maps having the same distribution of values and spatial autocorrelation, but different association with neuroanatomy [86]. Our results show that some cognitive topographies are more susceptible than others to facilitation via receptor-informed stimulation, whereas others exhibit little benefit (Fig. 5).

**Figure 5.**
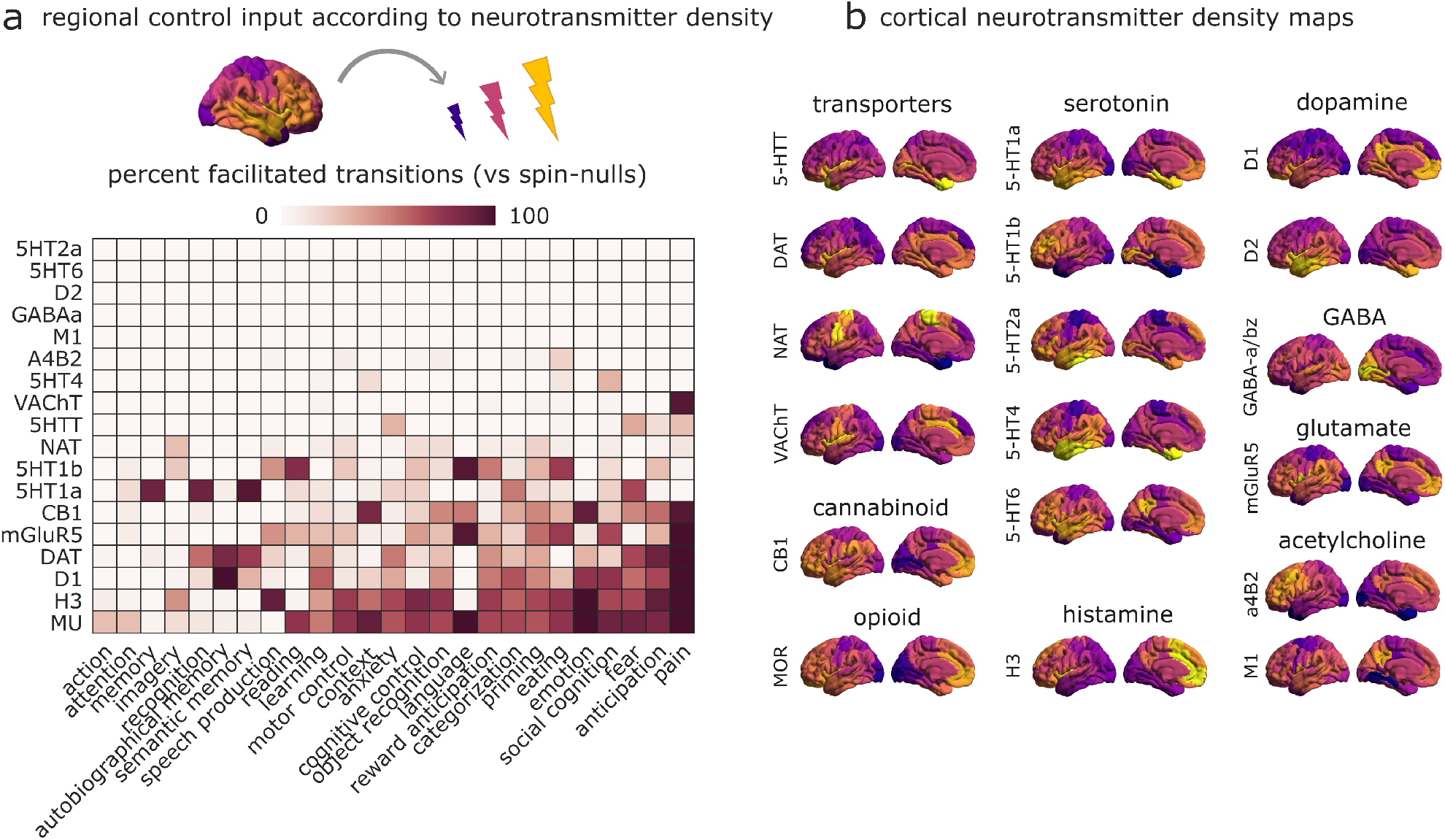
Modelling how neurotransmitter systems can reshape transition. **(a)** The effect of engaging each neurotransmitter receptor and transporter is modelled by changing the control input provided by each region in proportion to its density of receptor/transporter expression, measured by in vivo PET. Heatmap shows how each receptor/transporter reshapes the average cost of reaching a given cognitive brain state from all other states, as a percentage of transitions to each state that are facilitated, when compared against a null distribution of randomly rotated maps with preserved spatial autocorrelation and the same receptor/transporter density levels, but occurring at different neuroanatomical locations. **(b)** The empirical spatial distribution of each receptor and transporter is shown on the cortical surface.

Additionally, we find differences between receptors in terms of their propensity to facilitate transitions, over and above the mere effect of increased input (i.e., performing better than randomly rotated counterparts). Specifically, the dopamine transporter and D_1_ receptor, the mu-opioid receptor, and the histamine H_3_ receptor maps performed best (Fig. 5). Our computational framework identifies these receptors and transporters as those whose neuroanatomical distribution is mostly suited to facilitate transitions towards a variety of cognitive topographies.

### Replication and validation

We repeated our analyses using different parameter settings for network control theory; different reconstructions of the human connectome (with a different parcellation, and also in a separate DWI dataset); and a different way of defining cognitive topographies (based on the expert-curated BrainMap database [39, 76]). The *Supplementary Information* shows results with different implementations of the controllability framework [70]: we consider the time horizon *T* for control, adjacency matrix normalisation factor *c*, and the effect of normalising each map to unit Euclidean norm (Fig. S5, S6, S7, S8 and Tables S4, S5). We also show that our results can be replicated using a different reconstruction of the empirical human connectome, obtained from Diffusion Spectrum Imaging (Lausanne consensus dataset; [151]); and using a functional rather than anatomical parcellation of the human cerebral cortex to define network nodes (Schaefer-100 [120]) (Fig. S9).

Using the Lausanne consensus connectome, we see that the group-matrices of transition energy obtained using the HCP and Lausanne consensus connectomes are positively correlated (Spearman’s *r* = .99, *p* < 0.001). We also confirm that transitions between cognitive topographies are more energy-efficient on the human connectome than on degree-preserving nulls (*p* = 0.003). Although we do not find a significant difference between the empirical human connectome, and the distribution of null networks preserving both degree and wiring cost (*p* = 0.266), the superior energy-efficiency of the human connectome is confirmed at the single-subject level (Fig. S10a,b and Table S2). Both group-level and subject-level results are also replicated with network nodes defined by the Schaefer-100 atlas (Fig. S10c,d and Table S3).

The Lausanne dataset recapitulates the results about simulating pharmacological intervention (Fig. S12a). For Schaefer-parcellated data, we also observe a prominent role of D_1_ receptor and dopamine transporters, as before, but also acetylcholine and noradrenaline transporters (Fig. S12b). Lausanne dataset results are also consistent with HCP results in terms of the impact of cortical abnormality patterns (note that this analysis could not be repeated with the Schaefer-100 atlas, since ENIGMA data are only available in a single parcellation) (Fig. S11).

Finally, we show that analogous results can be obtained if instead of the automated NeuroSynth meta-analytic engine, we derive 66 cognitive topographies from BrainMap, an expert-curated database of published voxel coordinates from neuroimaging studies that are significantly activated or deactivated during tasks [39, 76]. As with NeuroSynth, we observe asymmetry of transitions, such that targets exhibit significantly greater variability in transition energy (mean = 8.54 × 10^4^, SD = 6.46 × 10^3^) than sources (mean = 4.27 × 10^4^, SD = 9.07 × 10^3^; *t*(130) = 31.21, *p* < 0.001, Cohen’s *d* = 5.40) (Fig. S13). Likewise we find that, among the BrainMap terms that have been assigned to one of the six cognitive ontology domains from [7], memory-related terms exhibit higher median value of asymmetry than would be expected by chance (*p* = 0.043). We also replicate the result that transitions between cognitive topographies require significantly less energy on the human connectome than on degree-preserving and degree-and cost-preserving null networks, with the geometry-preserving nulls being significantly closer to the human connectome than degree-preserving ones (all *p* < 0.001), Fig. S13).

## DISCUSSION

In the present report, we investigated how the brain’s network architecture shapes its capacity to transition between behaviourally defined cognitive topographies. We also systematically examined how transition between cognitive topographies could be reshaped by disease or pharmacological intervention. By taking into account network structure, functional activation and chemoarchitecture, our results provide a first step towards designing interventions that selectively manipulate cognitively relevant activation.

Up to this point, a comprehensive “look-up table” charting the transitions between cognitively relevant brain states has remained elusive. The present approach permits exploration of the full range of possible transitions between experimentally defined brain states with a cognitive interpretation. In this sense, our work provides a large-scale generalisation of recent advances [16, 68] that defined brain states in terms of *β* maps from a task-based fMRI contrast, or electrocorticography signal power associated with memory task performance [134]. Importantly, we have also expanded the network control framework to include naturalistic, empirically defined forms of control input, such as of receptor density (as could be exogenously engaged by pharmacological agents) and cortical abnormalities (as could arise endogenously from pathology).

We find that transitions between different cognitive topographies are not symmetric, but rather directional. Namely, some cognitively relevant brain states are substantially harder to reach than others, regardless of start state. These observations are consistent with the notion that state-to-state transition cost can perhaps be framed as cognitive demand [92]: in particular, transitioning to a more cognitively demanding 2-back task requires more control energy than transition to an easier task [16]. However, “cognitive demand” is a multifaceted construct with a variety of distinct possible operationalisations [47]: it remains to be determined which of these different interpretations is best aligned with network control energy. This endeavour will be facilitated by the approach introduced here, which enables the computational assessment of transition costs between any number of experimentally defined tasks from the literature.

Consistent with our work, a recent report identified a transition asymmetry between artificial “bottom-up” and “top-down” states, defined as recruiting different portions of the cytoarchitectonic sensory-fugal axis [106], finding that top-down states (i.e., involving a greater proportion of higher-order cortices) are more demanding. Indeed, we find that association of the target state with the putative unimodal-transmodal functional hierarchy is a predictor of transition cost. Moreover, the variability in ease-of-transition that we observe highlights cognitive topographies related to language and especially memory among those with the greatest asymmetry in transition cost. Both of these domains emerge gradually over human development [30, 100], and both language and autobiographical memory have long been argued (though not without controversy) to be “uniquely human” [84, 123, 136, 144].

Although there is variability among cognitive topographies in terms of transition cost, the wiring of the human connectome generally facilitates more efficient transitions than alternative topologies [73, 132, 139, 150]. Specifically, the human connectome enables transitions at lower cost compared to randomly rewired nulls that preserve degree sequence, suggesting that this efficiency is imparted by network topology, rather than low-level features such as the density and degree. Importantly, this efficiency can be partly attributed to the geometry of wiring lengths: when this was accounted for in our geometry-preserving null models, the connectome’s advantage was substantially diminished. Our work contributes to a growing appreciation for how network topology and geometry shape efficient communication [12, 19, 66, 133] and brain function [2, 96, 114, 138].

We also identified several factors that contribute to transition costs. Our results pertaining to Euclidean distance predicting transition costs are in line with those of Karrer and colleagues [70] and Stiso and colleagues [134], who found a monotonic increase of both minimum and optimal control energy with increasing distance between initial and target states. In terms of the states themselves, the best architectural predictor of transition cost to a given cognitive topography is its network-based variance (Fig. S2). A distribution of values over a network’s nodes has high network-based variance if nodes with high values are relatively difficult to reach using a diffusion process [29]. Network control theory predicts that control energy will diffuse along the network’s paths [132]. Therefore, we can interpret our results as showing that if a state requires great activation at nodes that are difficult to reach via diffusion, the corresponding state will be harder to reach. Indeed, network-based variance is related to bi-directional communicability between nodes—a known predictor of transition energy between states [14, 51, 106]. In other words, if a desired pattern of activations has low divergence from the pattern of diffusion-based proximities between nodes (low network-based variance), then that pattern will be easier to reach through network control.

Finally, we systematically quantified alternative naturalistic forms of control, via neurotransmitter receptor engagement and disorder-related aberrations in cortical morphology. Disorder-associated patterns of cortical abnormalities reshape the brain’s capacity to support transitions in ways that are not uniform, but rather both disorder-specific and brain state-specific. Of note, we find that depression and especially ADHD, both of which involve widespread attentional deficits [71, 145], are characterised by overall transition costs that exceed what would be expected if the corresponding cortical abnormalities were spatially distributed in a random fashion. Other disorders (schizophrenia, epilepsy) exhibited the opposite pattern. This suggests that localized perturbations in the connectome can attenuate some types of state transitions and promote others, potentially providing a mechanistic link between regional anatomical changes and changes in cognitive capacity across disease categories. This observation is consistent with results in the healthy population, where increased controllability is advantageous for cognitive performance in some regions, but disadvantageous in others [91].

Our results pertaining to pathology are complementary to applications of network control theory that evaluated transitions between random states or intrinsic connectivity networks, based on patients’ reorganised connectomes [11, 14, 58]—including a recent report that temporal lobe epilepsy induces deficits in control energy that are predictive of metabolic deficits quantified by FDG-PET [58]. In contrast, here we used the healthy connectome and simulated the effect of altered control input using abnormal cortical thickness (a change in control inputs rather than controlled network), and assessed the cost of transitioning between cognitive topographies that were experimentally defined. Since disorders typically involve both regional and connectomic alterations, in the future we expect that combining the two approaches will provide an even more fine-grained characterisation of how disease reshapes the brain’s capacity to transition between brain states.

We observed a similar principle when control inputs were guided by empirically-derived receptor and transporter maps. We find that the more difficult cognitive topographies to reach also appear to be those that can most benefit from pharmacological intervention. Here dopamine transporters and D1 receptors, histamine H3 receptors, and mu-opioid receptors appear well-positioned to facilitate transitions. These results are consistent with the use of modafinil and methylphenidate as cognitive enhancers and to treat symptoms of ADHD: both drugs engage the dopaminergic system by blocking dopamine transporter as one of their main mechanisms of action [38, 46, 64, 65, 74, 83, 87, 95, 135, 146, 152–154, 157]. Likewise, H_3_-receptor antagonist drugs such as pitolisant are being evaluated for potential treatment of ADHD symptoms [37, 57, 105, 155].

Collectively, the predictions generated by our report highlight numerous potential clinical and non-clinical applications. Although preliminary, these results provide a first step towards designing protocols that selectively promote transitions to desired cognitive topographies in specific diseases. In addition, outcomes of this computational screening could be further tested *in vivo* by engaging different neurotransmitter systems through targeted pharmacological manipulations [16, 124], and evaluating the degree to which they facilitate switching between specific experimentally-defined cognitive topographies.

The present work should be interpreted with respect to several important methodological considerations. Network control theory models neural dynamics as noise-free, and under assumptions of linearity and time invariance [52, 70, 73]. Recent work has begun to introduce stochasticity in the network control framework for the brain, with promising results—though still within the context of linear systems [68]. Although the brain is a nonlinear system, it has been shown that non-linear dynamics can be locally approximated by linear dynamics [26, 62], including through the application of dynamic causal models [40, 41]. In fact, evidence suggests that linear models may even outperform nonlinear ones at the macroscopic scale of BOLD signals [103, 122]. Finally, the predictions of linear network control theory have found successful translation to nonlinear systems [97, 143], and even at predicting the effects of direct intracranial electrical stimulation in humans [134].

Additionally, we made several simplifying assumptions about the control input provided by each region based on its cortical thickness or receptor/transporter expression. We have treated summary statistics from NeuroSynth’s term-based meta-analysis as representing relative activation; we also acknowledge that the mapping of functional activation to psychological terms in NeuroSynth does not distinguish activations from deactivations [159]. However, we believe that our replication with cognitive topographies defined using Brain-Map [39] provides reassurance about the validity of our approach. Although the ENIGMA consortium provides datasets from large cohorts with standardised pipelines, ensuring robust results, the patient populations may exhibit co-morbidities and/or be undergoing treatment. In addition, and of particular relevance for the present modelling approach, the available maps do not directly reflect changes in tissue volume, but rather the effect size (magnitude of between-group difference) of patient-control statistical comparisons (though note that our use of spatial autocorrelation-preserving null models accounts for the mean and variance of each map; see *Methods).* Moreover, many more disorders, diseases, and conditions exist than the ones considered here. The same limitation applies to the PET data: the atlas of neurotransmitter receptors, though extensive, does not include all receptors. However, our computational workflow can readily be extended to accommodate new cognitive topographies, receptors, or disease maps of interest. Finally, here we did not consider the role of the subcortex, which is not present in ENIGMA disorder maps, and needs different treatment both in terms of spatial null models, and in terms of PET imaging. This approach has also been adopted in recent applications of network control theory to the human connectome [106, 107].

Overall, our approach based on network control theory provides a computational framework to evaluate the propensity of the connectome to support transitions between cognitively-relevant brain patterns. This framework lends itself to interrogating transitions between specific states of interest, and modelling the impact of global perturbations of the connectome, or regional cortical heterogeneities, or specific neurotransmitter systems. We anticipate that future work may combine different facets of this approach to evaluate *in silico* which potential pharmacological treatments may best address the specific cognitive difficulties associated with a given disorder or brain tissue lesion.

## METHODS

### Human structural connectome from Human Connectome Project

We used diffusion MRI (dMRI) data from the 100 unrelated subjects of the HCP 900 subjects data release [148]. All HCP scanning protocols were approved by the local Institutional Review Board at Washington University in St. Louis. The diffusion weighted imaging (DWI) acquisition protocol is covered in detail elsewhere [48]. The diffusion MRI scan was conducted on a Siemens 3T Skyra scanner using a 2D spin-echo single-shot multiband EPI sequence with a multi-band factor of 3 and monopolar gradient pulse. The spatial resolution was 1.25 mm isotropic. TR=5500 ms, TE=89.50ms. The b-values were 1000, 2000, and 3000 s/mm^2^. The total number of diffusion sampling directions was 90, 90, and 90 for each of the shells in addition to 6 b0 images. We used the version of the data made available in DSI Studio-compatible format at http://brain.labsolver.org/diffusion-mri-templates/hcp-842-hcp-1021 [160].

We adopted previously reported procedures to reconstruct the human connectome from DWI data. The minimally-preprocessed DWI HCP data [48] were corrected for eddy current and susceptibility artifact. DWI data were then reconstructed using q-space diffeomorphic reconstruction (QSDR [162]), as implemented in DSI Studio (www.dsi-studio.labsolver.org). QSDR is a model-free method that calculates the orientational distribution of the density of diffusing water in a standard space, to conserve the diffusible spins and preserve the continuity of fiber geometry for fiber tracking. QSDR first reconstructs diffusion-weighted images in native space and computes the quantitative anisotropy (QA) in each voxel. These QA values are used to warp the brain to a template QA volume in Montreal Neurological Institute (MNI) space using a nonlinear registration algorithm implemented in the statistical parametric mapping (SPM) software. A diffusion sampling length ratio of 2.5 was used, and the output resolution was 1 mm. A modified FACT algorithm [161] was then used to perform deterministic fiber tracking on the reconstructed data, with the following parameters [82]: angular cutoff of 55o, step size of 1.0 mm, minimum length of 10 mm, maximum length of 400 mm, spin density function smoothing of 0.0, and a QA threshold determined by DWI signal in the cerebrospinal fluid. Each of the streamlines generated was automatically screened for its termination location. A white matter mask was created by applying DSI Studio’s default anisotropy threshold (0.6 Otsu’s threshold) to the spin distribution function’s anisotropy values. The mask was used to eliminate streamlines with premature termination in the white matter region. Deterministic fiber tracking was performed until 1, 000, 000 streamlines were reconstructed for each individual.

For each individual, their structural connectome was reconstructed by drawing an edge between each pair of regions *i* and *j* from the Desikan-Killiany cortical atlas [27] if there were white matter tracts connecting the corresponding brain regions end-to-end; edge weights were quantified as the number of streamlines connecting each pair of regions, normalised by ROI distance and size.

A group-consensus matrix *A* across subjects was then obtained using the distance-dependent procedure of Betzel and colleagues, to mitigate concerns about inconsistencies in reconstruction of individual participants’ structural connectomes [13]. This approach seeks to preserve both the edge density and the prevalence and length distribution of inter- and intra-hemispheric edge length distribution of individual participants’ connectomes, and it is designed to produce a representative connectome [13, 96]. The final edge density was 27%, and the weight associated with each edge was computed as the mean non-zero weight across participants.

### Alternative structural connectome from Lausanne dataset

A total of *N* = 70 healthy participants (25 females, age 28.8 ± 8.9 years old) were scanned at the Lausanne University Hospital in a 3-Tesla MRI Scanner (Trio, Siemens Medical, Germany) using a 32-channel head coil [50]. Informed written consent was obtained for all participants in accordance with institutional guidelines and the protocol was approved by the Ethics Committee of Clinical Research of the Faculty of Biology and Medicine, University of Lausanne, Switzerland. The protocol included (1) a magnetization-prepared rapid acquisition gradient echo (MPRAGE) sequence sensitive to white/gray matter contrast (1 mm in-plane resolution, 1.2 mm slice thickness), and (2) a diffusion spectrum imaging (DSI) sequence (128 diffusion-weighted volumes and a single b0 volume, maximum b-value 8 000 s/mm^2^, 2.2 × 2.2 × 3.0 mm voxel size).

Structural connectomes were reconstructed for individual participants using deterministic streamline tractography and divided according to the Desikan-Killiany grey matter parcellation. White matter and grey matter were segmented from the MPRAGE volumes using the FreeSurfer version 5.0.0 open-source package, whereas DSI data preprocessing was implemented with tools from the Connectome Mapper open-source software, initiating 32 streamline propagations per diffusion direction for each white matter voxel. Structural connectivity was defined as streamline density between node pairs, i.e., the number of streamlines between two regions normalized by the mean length of the streamlines and the mean surface area of the regions, following previous work with these data [53, 151].

### Network control energy

Network control theory models the brain as a linear, time-invariant control system [52]. In the general context of linear control theory, the evolutionary dynamics of the state *x*(*t*) is formulated as an equation relating the first-order derivative of the state *x*(*t*), 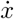, to the state variable *x* itself, and the control input. For this system, given the initial and target states, the control trajectory moving from the initial to target states is determined by the interaction matrix *A*, the input matrix *B*, and the control input *u*(*t*), in the form of

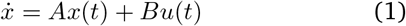

The state interaction matrix *A* characterizes the relationships between system elements, determining how the control system moves from the current state to the future state. The structural connectivity matrix *A* serves as a linear operator that maps each state, *x*, to the rate of change of that state. This linear transformation can be described in terms of the evolutionary modes of the system consisting of the *N* eigenvectors of *A* and their associated eigenvalues.

The matrix *A* is then normalised to avoid infinite growth of the system over time:

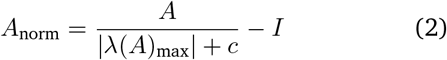

Here, *I* denotes the identity matrix of size *N* × *N*, and |λ(*A*)_max_| denotes the largest eigenvalue of the system. To normalize the system, we must specify the parameter *c*, which determines the rate of stabilization of the system. Here we use *c* = 0 for our main analyses, such that the system approaches its largest mode over time. We also report results for *c* = 0.01 × |λ(*A*)_max_|, whereby all modes decay, and the system goes to zero over time. The control input matrix B denotes the location of control nodes on which we place the input energy. If we control all brain regions, B corresponds to the *N* × *N* identity matrix with ones on the diagonal and zeros elsewhere. If we control only a single brain region *i, B* reduces to a single *N* × *N* diagonal matrix with a one in the *i*^th^ element of the diagonal, and zeros elsewhere. The control input *u*(*t*) denotes the amount of energy injected into each control node at each time point *t*. Intuitively, *u*(*t*) can be summarized over time to represent the total energy consumption during transition from an initial state to a final state. In the brain control analysis framework, a state refers to a vector *x*(*t*) of *N* elements, which encodes the neurophysiological activity map across the whole brain. In the current work, *x*(*t*) is the meta-analytic activation of each region associated with each cognitively relevant term, aggregated over studies in the NeuroSynth database.

This computational approach allows us to compute the transition energy as the optimal energy required to transition between each pair of cognitive topographies in finite time.

To explore the energetic efficiency of the structural brain network in facilitating the transition between cognitive topographies, we adopted the optimal control framework to estimate the control energy required to optimally steer the brain through these state transitions [14, 51, 58]. Optimality is defined in terms of jointly minimising the combination of both the length of the transition trajectory from an initial source state (*x*(0) = *x*_0_) to the final target state (*x*(*T*) = *x_T_*) over the time horizon *T* (to avoid spurious, unrealistically long trajectories), and the required unique control input *u*\*(*t*) summarised over the length of this trajectory:

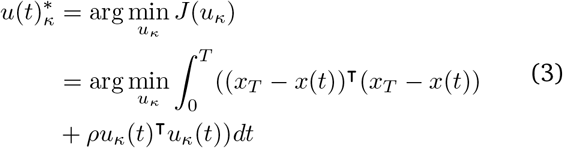

where (*x_T_* − *x*(*t*))^⊤^(*x_T_* – *x*(*t*)) is the distance between the state at time *t* and the final state *x_T_*, *T* is the finite amount of time given to reach the final state, and *ρ* is the relative weighting between the cost associated with the length of the transition trajectory and the input control energy. The equation is solved using forward integration. 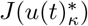 is the cost function defined to find the unique optimal control input 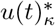. Here, following common practice, we set *ρ* equal to 1, corresponding to equal weighting [14, 70]. We set *T* =1 [14], but we also report results with *T* = 3 [70] (Fig. S5, S6, S7, S8).

### Cognitive topographies from NeuroSynth

Continuous measures of the association between voxels and cognitive categories were obtained from NeuroSynth, an automated term-based meta-analytic tool that synthesizes results from more than 15 000 published fMRI studies by searching for high-frequency key words (such as “pain” and “attention” terms) that are systematically mentioned in the papers alongside fMRI voxel coordinates (https://github.com/neurosynth/neurosynth, using the volumetric association test maps [159]). This measure of association strength is the tendency that a given term is reported in the functional task-based neuroimaging study if there is activation observed at a given voxel. Note that the tool does not distinguish between areas that are activated or deactivated in relation to the term of interest, nor the degree of activation, only that certain brain areas are frequently reported in conjunction with certain words. Although more than a thousand terms are catalogued in the NeuroSynth engine, we refine our analysis by focusing on cognitive function and therefore we limit the terms of interest to cognitive and behavioural terms. These terms were selected from the Cognitive Atlas, a public ontology of cognitive science [109], which includes a comprehensive list of neurocognitive terms and has been previously used in conjunction with NeuroSynth [54]. This approach totaled to t = 123 terms, ranging from umbrella terms (“attention”, “emotion”) to specific cognitive processes (“visual attention”, “episodic memory”), behaviours (“eating”, “sleep”), and emotional states (“fear”, “anxiety”). The probabilistic measure reported by Neurosynth can be interpreted as a quantitative representation of how regional fluctuations in activity are related to psychological processes.

### Alternative cognitive topographies from BrainMap

Whereas NeuroSynth is an automated tool, Brain-Map is an expert-curated repository: it includes the brain coordinates that are significantly activated during thousands of different experiments from published neuroimaging studies [39, 76]. As a result, NeuroSynth terms and BrainMap behavioural domains differ considerably. Here, we used maps pertaining to 66 unique behavioural domains (the same as in [54]), obtained from 8,703 experiments. Experiments conducted on unhealthy participants were excluded, as well as experiments without a defined behavioural domain.

### Network null models

We used two different network null models to disambiguate the role of connectome topology and geometric embedding in shaping control energy [150]. The first null model is the well-known Maslov-Sneppen degree-preserving rewired network, whereby edges are swapped so as to randomise the topology while preserving the exact binary degree of each node (degree sequence), and the overall distribution of edge weights [88]. As a second, more stringent null model, we adopted a null model that in addition to preserving exactly the same degree sequence and exactly the same edge weight distribution as the original network, also approximately preserves the original network’s edge length distribution (based on Euclidean distance between regions), and the weight-length relationship [12].

For each null model, we generated a population of 500 null networks starting from the empirical connectome, and computed the control energy between each pair of cognitive brain states from NeuroSynth, as done for the empirical connectome. We compared the overall control energy between all possible states obtained from the empirical connectome and from the distribution of null instances, by obtaining a *z*-score.

### Spatial null models

To evaluate the role of regional neuroanatomical features, we implemented a permutation-based null model, termed spin test [86, 150]. For each map, parcel coordinates were projected onto the spherical surface and then randomly rotated and original parcels were reassigned the value of the closest rotated parcel (10 000 repetitions) [151]. In addition to preserving the distribution of cortical values, this null model also preserves the spatial autocorrelation present in the data.

### Predictors of transition energy

We characterised each cognitive brain state from NeuroSynth in terms of its relationship with several well-known graph-theoretic properties of the structural connectome. From the consensus connectome we computed the binary and weighted degree (also known as node strength) of each region. We also computed the participation coefficient of each region, based on the modular assignment of each region to the well-known intrinsic connectivity networks [163]. We computed the Spearman correlation between each of these vectors, and each of the 123 brain maps from NeuroSynth. Additionally, we also computed the correlation between each NeuroSynth map, and the principal gradient of variation in functional connectivity [85], believed to reflect the hierarchical organisation of cortical information-processing.

### Network-based variance

For each NeuroSynth map, we also computed as an additional predictor a recently developed measure termed “network variance” [29]. The traditional notion of variance of a distribution (sum of squared differences from the mean) assumes that observations are independent. However, this assumption is almost invariably violated in the case of distributions on a graph, where the graph’s nodes (whose values correspond to the distribution’s observations) are connected to each other, genereating dependencies. The notion of spatial autocorrelation [86] can be cast as a special case of this situation, whereby the graph connecting nodes is the graph of spatial distances between them (e.g., Euclidean distance). Devriendt and colleagues provided the network variance as a generalization of variance to distributions on a graph:

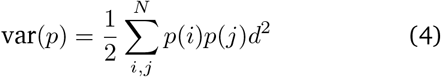

In other words, a distribution on a graph has high network variance if most of the mass (here: activity) is concentrated at nodes that are poorly connected with the rest of the network. This relies on defining a suitable measure of distance on a graph. Devriendt and colleagues noted that the geodesic distance (length of the shortest path between two nodes) may be a suitable candidate, but recommended using the effective resistance instead [28, 29]. Like geodesic distance, the effective resistance is predicated on the length of the paths between a pair of nodes. However, unlike the geodesic distance, effective resistance does not only consider the shortest path between two nodes, but rather it takes into account paths of all length along the graph, such that two nodes are less distant the more paths exist between them, thereby reflecting the full topology of the network. The resistance distance *ω_ij_* between nodes *i* and *j* is large when nodes *i* and *j* are not well connected in the network, such that only few, long paths connect them, resulting in a long time for a random walker to reach one node from another, whereas a small *ω_ij_* means that they are well connected through many, predominantly short paths *i* and *j* [28]. Up to a constant, the effective resistance can be computed as the “commute time”: the mean time it takes a random walker to go from node *i* to node *j* and back, for all pairs of nodes *i* and *j* [21]. Concretely, effective resistance on a graph is computed as

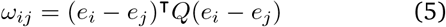

where the unit vectors have entries (*e_i_*)*k* = 1 if *k* = *i* and zero otherwise, and where *Q* is the Moore-Penrose pseudoinverse of the graph’s Laplacian matrix.

For each NeuroSynth map, we computed its network variance using as distance measure the effective resistance on the consensus connectome. Since the network variance requires the distribution on the graph’s nodes to be positive and sum to 1, each map’s values were rescaled so that the minimum was 0, and then divided by their sum. We then used these characterisations of the NeuroSynth maps (correlation with connectome graph-theoretic properties; correlation with the cortical hierarchy; and network variance) as predictors against the average energy required to transition to each cognitive brain state. We performed multiple partial correlations, using each characterisation in turn as predictor (after partialling out the effects of mean and traditional variance of each NeuroSynth map).

### Dominance analysis

As an alternative approach, to consider all predictors together and evaluate their respective contributions, we performed a dominance analysis with all five predictors. Dominance analysis seeks to determine the relative contribution (“dominance” of each independent variable to the overall fit (adjusted *R*^2^)) of the multiple linear regression model (https://github.com/dominance-analysis/dominance-analysis) [4]. This is done by fitting the same regression model on every combination of predictors (2^*p*^ – 1 submodels for a model with *p* predictors). Total dominance is defined as the average of the relative increase in *R*^2^ when adding a single predictor of interest to a submodel, across all 2^*p*^ – 1 submodels. The sum of the dominance of all input variables is equal to the total adjusted *R*^2^ of the complete model, making the percentage of relative importance an intuitive method that partitions the total effect size across predictors. Therefore, unlike other methods of assessing predictor importance, such as methods based on regression coefficients or univariate correlations, dominance analysis accounts for predictor–predictor interactions and is interpretable.

### Disease-associated patterns of cortical thickness abnormality from the ENIGMA database

Spatial maps of cortical thickness were collected for the available 11 neurological, neurodevelopmental, and psychiatric disorders from the ENIGMA (Enhancing Neuroimaging Genetics through Meta-Analysis) consortium [56, 141, 142] and the Enigma toolbox and recent related publications (https://github.com/MICA-MNI/ENIGMA) [77]: 22q11.2 deletion syndrome [137], attention-deficit/hyperactivity disorder [63], autism spectrum disorder [149], idiopathic generalized epilepsy [156], right temporal lobe epilepsy [156], left temporal lobe epilepsy [156], depression [121], obsessive-compulsive disorder [15], schizophrenia [147], bipolar disorder [59], and Parkinson’s disease [75]. The ENIGMA consortium is a data-sharing initiative that relies on standardized image acquisition and processing pipelines, such that disorder maps are comparable [141]. Altogether, over 17 000 patients were scanned across the eleven disorders, against almost 22 000 controls. The values for each map are z-scored effect sizes (Cohen’s *d*) of cortical thickness in patient populations versus healthy controls. Imaging and processing protocols can be found at http://enigma.ini.usc.edu/protocols/.

For every brain region, we constructed an 11-element vector of disorder abnormality, where each element represents a disorder’s cortical abnormality at the region. These values were then added to the *B* matrix of uniform control inputs, to provide regional heterogeneity. This approach is motivated by the expectation that regions with decreased thickness should have lower capacity to exert control inputs, and vice-versa. Recent work adopted a similar approach to model the regional control input provided at each region, in terms of the regional density of receptor expression, such that regions expressing the receptor to a greater extent are understood to exert greater control input [124]. The largest increase in cortical thickness across all disorders is 0.87 (expressed in terms of Cohen’s d), whereas the largest decrease is 0.59. Thus, across all diseases, the entries in the input matrix B were always positive, bound between 0.41 and 1.87. Since the distributions of cortical abnormality associated with the various ENIGMA diseases and disorders are different, changing the distribution of control inputs also changes the overall amount of control energy that is being injected into the system (in some cases leading to an overall increase, or an overall decrease), and consequently the control energy that is required to transition between brain states. Therefore, an appropriate null model to evaluate the effects of disease-associated patterns of grey matter abnormality on brain state transitions should preserve the overall spatial distribution of control inputs associated with each map, while changing the spatial location. Rather than simply randomising the distribution of cortical thickness abnormalities, we opted to adopt the spin-based null model, which also preserves the spatial autocorrelation present in the data [86, 150]. We then obtained a z-score of the control energy required to transition between each pair of states (here considering only the reduced set of 25 states, to reduce computational burden), for the empirical ENIGMA map against the distribution of spin-null maps. Thus, a positive z-score indicates that the empirical pattern of cortical thickness abnormality associated with a disease is more energetically demanding than would be expected if the abnormalities were occurring at random on the cortex; whereas a negative z-score indicates that the pattern requires less energy than if the abnormalities were occurring at random (but with equivalent spatial autocorrelation).

### Receptor maps from Positron Emission Tomography

Receptor densities were estimated using PET tracer studies for a total of 18 receptors and transporters, across 9 neurotransmitter systems, recently made available by Hansen and colleagues at https://github.com/netneurolab/hansen_receptors [55]. These include dopamine (D_1_ [67], D_2_ [117, 125, 128, 164], DAT [35], norepinephrine (NET [9, 31, 79, 116], serotonin (5-HT_1A_ [119], 5-HT_1B_ [42, 89, 98, 108, 118, 119], 5-HT_2A_ [10], 5-HT_4_ [10], 5-HT_6_ [111, 112] 5-HTT [10]), acetylcholine (*α*_4_*β*_2_ [5, 60], M_1_ [99], VAChT [1, 8], glutamate (mGluR_5_ [34, 127] NMDA [44, 45], GABA (GABAA [104])), histamine (H_3_ [43]), cannabinoid (CB_1_ [33, 101, 102, 113]) and opioid (MOR [69]). Volumetric PET images were registered to the MNI-ICBM 152 nonlinear 2009 (version c, asymmetric) template, averaged across participants within each study, then parcellated and receptors/transporters with more than one mean image of the same tracer (5-HT_1_B, D_2_, VAChT) were combined using a weighted average [55].

For the control energy analysis, each PET map was scaled between 0 and 1, and its regional values were added to the B matrix, following recent work [124]. Since the PET distributions are different, changing the distribution of control inputs also changes the overall amount of control energy that is being injected into the system, and consequently the control energy that is required to transition between brain states. Therefore, an appropriate null model to evaluate which receptors are especially well-poised to facilitate brain state transitions, in terms of their spatial location, should preserve the overall distribution of control inputs associated with each receptor, while changing the spatial location. Rather than simply randomising the distribution of receptor densities, we opted to adopt the more stringent spin test null [86, 150]. In addition to preserving the distribution of regional receptor densities, this null model also preserves the spatial autocorrelation present in the data. We then evaluated how often (as a percentage) a transition between two given brain states was found to require less control energy when using an empirical receptor map to determine the control inputs, than when using a spin-randomised version of the same map. Due to the computationally intensive nature of this procedure (10 000 repetitions for each of 18 PET maps), we only considered transitions between the reduced set of 25 brain states, instead of the whole 123.

## Acknowledgments

We thank Vincent Bazinet, Eric Ceballos, Filip Milisav, Laura Suarez, Zhen-Qi Liu, and Moohebat Pourmajidian for helpful discussion. AIL is supported by the Molson Neuro-Engineering Fellowship and FRQNT Strategic Clusters Program (2020-RS4-265502 - Centre UNIQUE - Union Neuroscience & Artificial Intelligence - Quebec) via the UNIQUE Neuro-AI Excellence Scholarship. SPS is supported by the National Science Foundation Graduate Research Fellowship (Grant No. DGE-1650441). JYH is supported by the Helmholtz International BigBrain Analytics & Learning Laboratory, the Natural Sciences and Engineering Research Council of Canada, and Fonds de reserches de Québec. DB is supported by the Brain Canada Foundation, through the Canada Brain Research Fund, with the financial support of Health Canada, National Institutes of Health (grants no. NIH R01 AG068563A and NIH R01 R01DA053301-01A1 to DB), the Canadian Institute of Health Research (grants no. CIHR 438531 and CIHR 470425 to DB), the Healthy Brains Healthy Lives initiative (Canada First Research Excellence fund), Google (Research Award, Teaching Award to DB) and by the CIFAR Artificial Intelligence Chairs programme (Canada Institute for Advanced Research to DB). AK is supported by the National Institutes of Health (RF1MH123232, R01NS102646 and R21NS104634). BM acknowledges support from the Natural Sciences and Engineering Research Council of Canada (NSERC), Canadian Institutes of Health Research (CIHR), Brain Canada Foundation Future Leaders Fund, the Canada Research Chairs Program, the Michael J. Fox Foundation, and the Healthy Brains for Healthy Lives initiative. Human data were provided by the Human Connectome Project, WU-Minn Consortium (1U54MH091657; Principal Investigators David Van Essen and Kamil Ugurbil) funded by the 16 National Institutes of Health (NIH) institutes and centers that support the NIH Blueprint for Neuroscience Research, and by the McDonnell Center for Systems Neuroscience at Washing-ton University.

## Supplementary Materials

**Figure S1.**
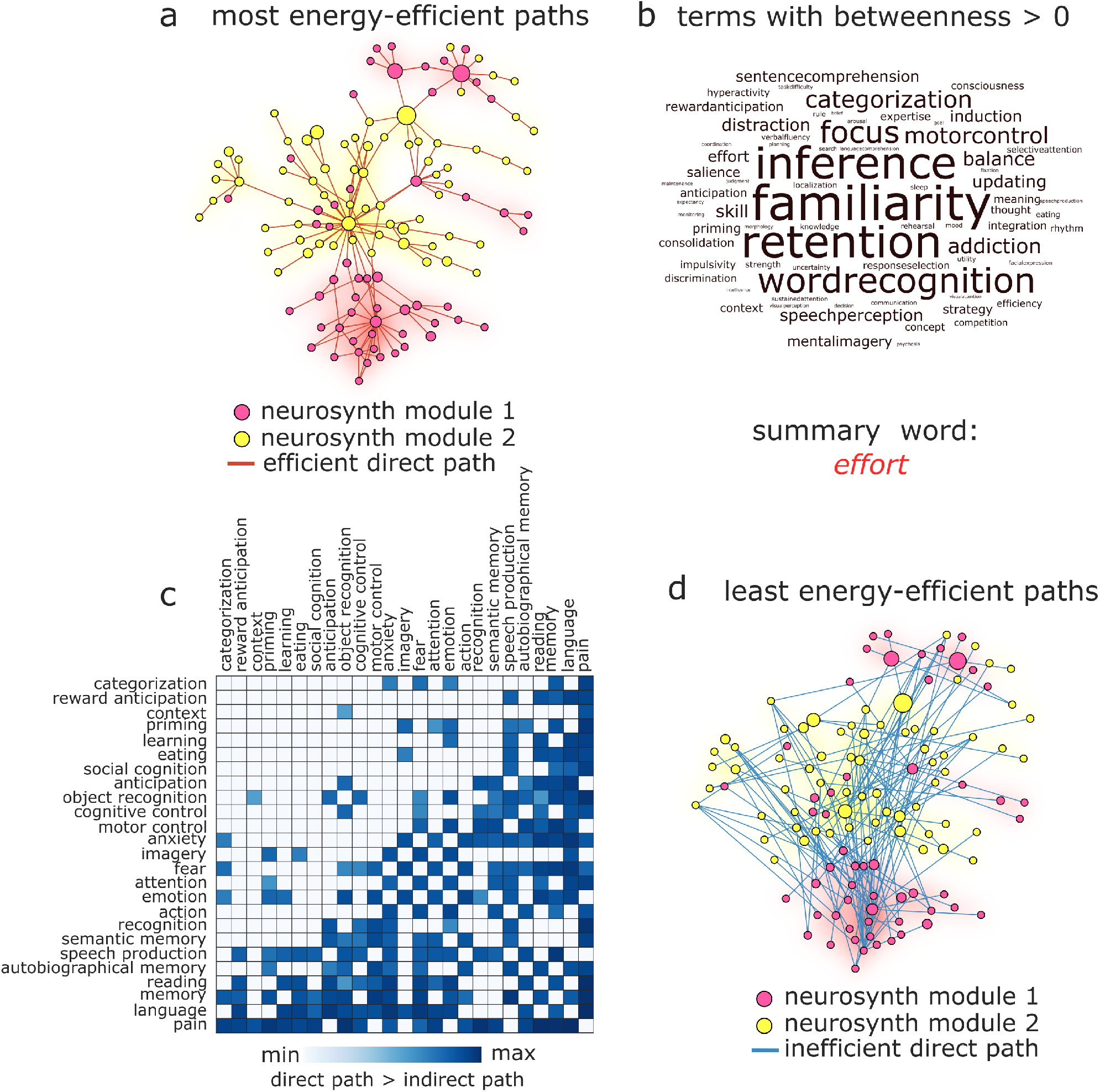
Efficient and inefficient paths between cognitive topographies. **(a)** Representation of the control energy matrix as a weighted network, showing the transitions (edges) between cognitive topographies (nodes) that require the least energy (for display purposes, only 10% of connections are shown). Nodes are colored according to their membership of the two communities into which cognitive topographies cluster. Cognitive topographies that act as intermediaries for the most efficient transition between two other cognitive topographies correspond to nodes that have non-zero betweenness centrality. Node size reflects the betweenness centrality of each node (taking into account all paths). **(b)** The NeuroSynth terms corresponding to cognitive topographies with non-zero betweenness centrality in the network representation in (a); size reflects the betweenness centrality of the corresponding nodes. To identify the term that most summarises all others, we represent each term as a high-dimensional vector in semantic space using *word2vec* [94], and we measure their similarity using cosine similarity between these vector representations. We find that “effort” has the highest mean cosine similarity with the vector representations of all other high-betweenness terms. **(c)** Matrix of the transition cost between each pair of cognitive topographies in the reduced set, showing the difference in control energy between the direct and least-expensive paths; the value of each non-empty cell indicates the energy premium incurred by taking the direct path between two cognitive topographies. **(d)** The most costly direct paths between cognitive topographies (only 10% shown, for display purposes). Node size, colour, and position are the same as in (a).

**Figure S2.**
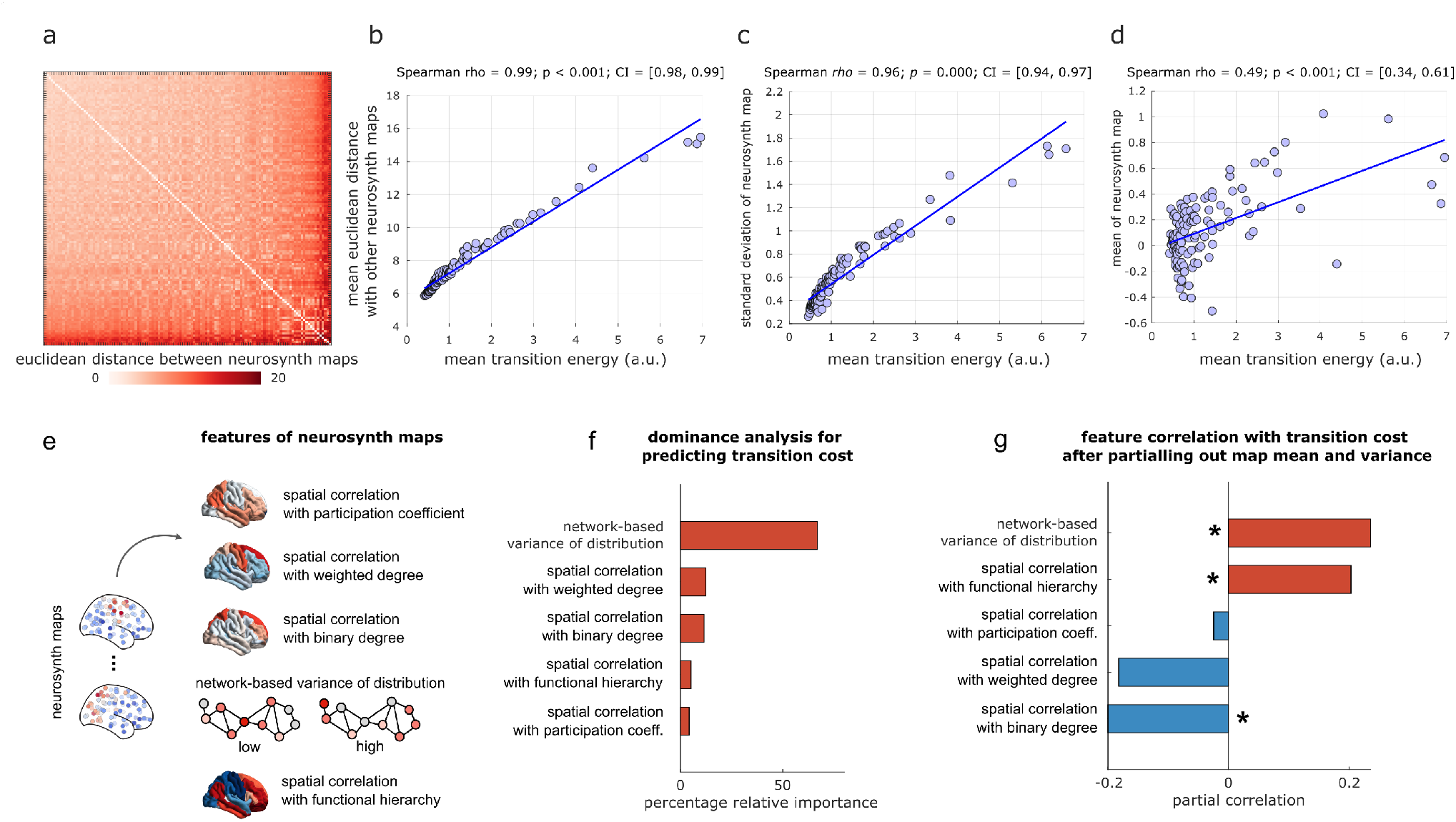
Predictors of transition energy. **(a)** Euclidean distance between the vectors corresponding to each NeuroSynth map. **(b)** Transition energy to a given cognitive topography (averaged over all starting states) correlates with the mean Euclidean distance between its corresponding NeuroSynth map and all others. **(c)** Transition energy to a given cognitive topography (averaged over all starting states) correlates with the standard deviation of its NeuroSynth map. **(d)** Transition energy to a given cognitive topography (averaged over all starting states) correlates with the mean of its NeuroSynth map. **(e)** Mean and variance of a NeuroSynth map are coarse descriptions that do not account for neuroanatomy. As neuroanatomically-grounded predictors of transition energy, we consider the following features of each NeuroSynth map: (i) the map’s spatial alignment with the regional distribution of participation coefficients, treating the structural connectome as a network; (ii) the map’s spatial alignment with the regional distribution of weighted node degree, treating the structural connectome as a network; (iii) the map’s spatial alignment with the regional distribution of binary node degree, treating the structural connectome as a network; (iv) a recently developed measure of variance for distributions over a network [28, 29]: unlike the usual measure of variance, which assumes independent data-points and is agnostic to their spatial location, this measure takes into account the relationships between observations. Specifically, variance of a distribution over a network is low, if high values occur at nodes that are easy to reach using a diffusion process along network paths of all length. Conversely, if the majority of high values occurs at nodes that are difficult to reach using diffusion, then the distribution will have high network-based variance. This is especially relevant because network control theory operationalises control inputs as spreading by diffusing over the network. (v) The map’s spatial alignment with the principal gradient of functional connectivity (unimodal-transmodal hierarchy) [85]. **(f)** Bar plot shows the relative dominance of each predictor as obtained from dominance analysis [4]. Dominance analysis distributes the fit of the model across predictors such that the contribution of each predictor can be assessed and compared to other predictors, reflecting the proportion of the variance jointly explained by all predictors, that can be attributed to each predictor. **(g)** Bar plot shows the partial correlation between each predictor and the average cost to transition to a given brain state, after controlling for the effects of brain state mean and variance.

**Figure S3.**
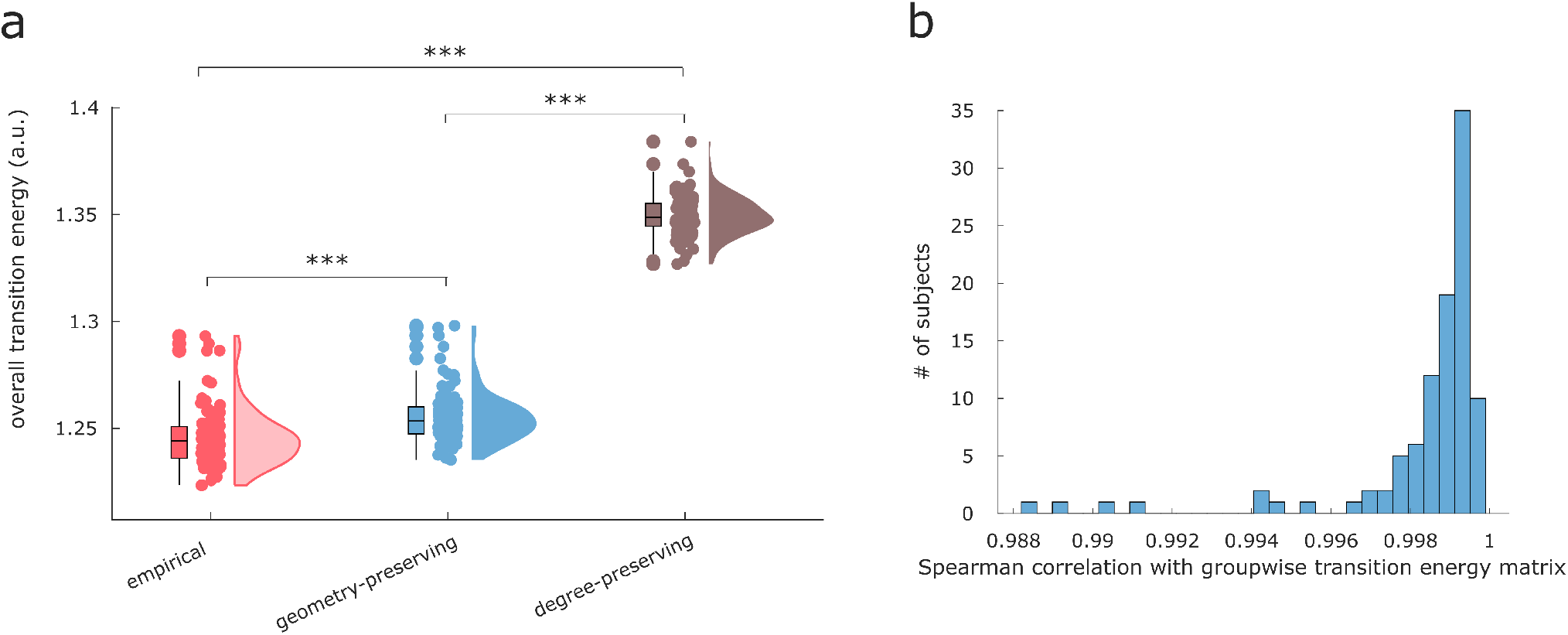
Replication at the single-subject level. **(a)** Overall transition energy (averaged across all transitions), for each subject and for the corresponding degree- and cost-preserving nulls. **(b)** Histogram of the correlation between each subject’s matrix of transition energies, and the matrix of transition energies obtained from the group-wise consensus structural connectome.

**Figure S4.**
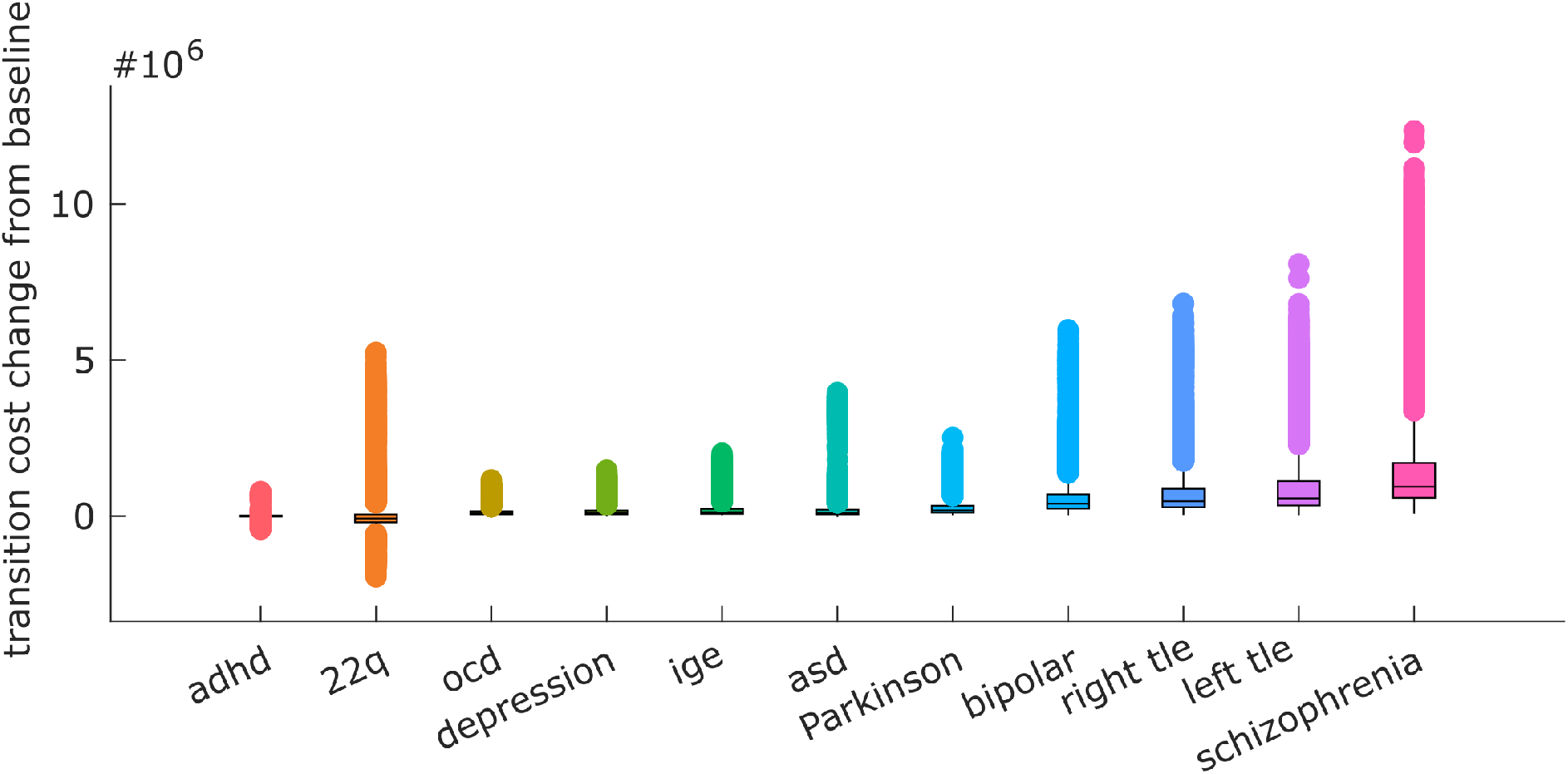
Overall transition energy when applying control inputs according to cortical abnormality maps. Each data-point represents the energy to transition to one target state, averaging across all source states.

**Figure S5.**
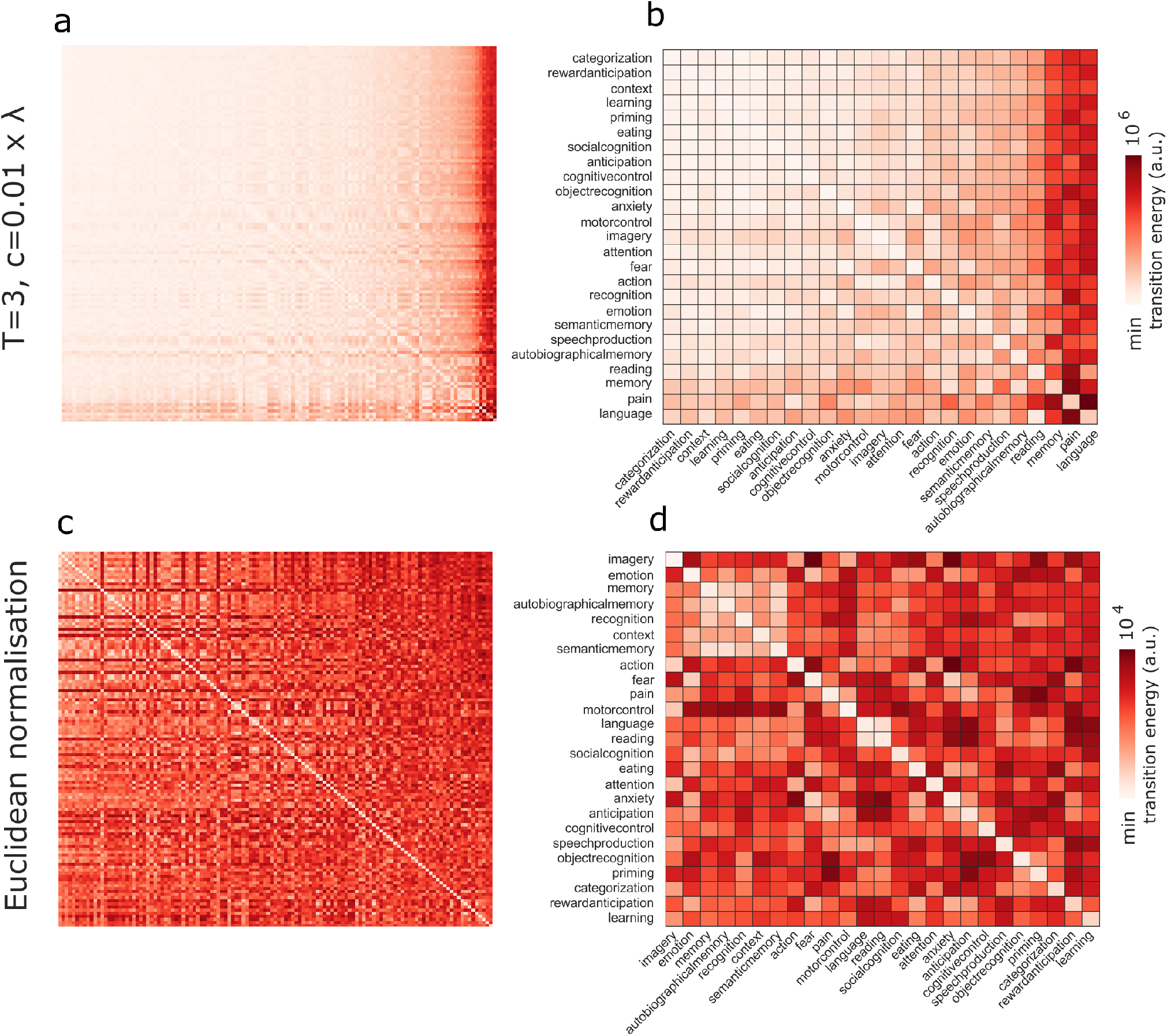
Transition energies for alternative operationalisations of network control theory. **(a,b)** Transition energy between each pair of 123 cognitive topographies from NeuroSynth (a) and the reduced set of 25 NeuroSynth terms (b), for network control with time horizon *T* = 3 and network normalisation factor *c* = 0.01 × |λ(*A*)_max_|. Rows indicate source states, columns indicate target states. **(c,d)** Transition energy between each pair of 123 cognitive topographies from NeuroSynth (c) and the reduced set of 25 NeuroSynth terms (d), for network control with all NeuroSynth maps normalised to unit Euclidean norm.

**Figure S6.**
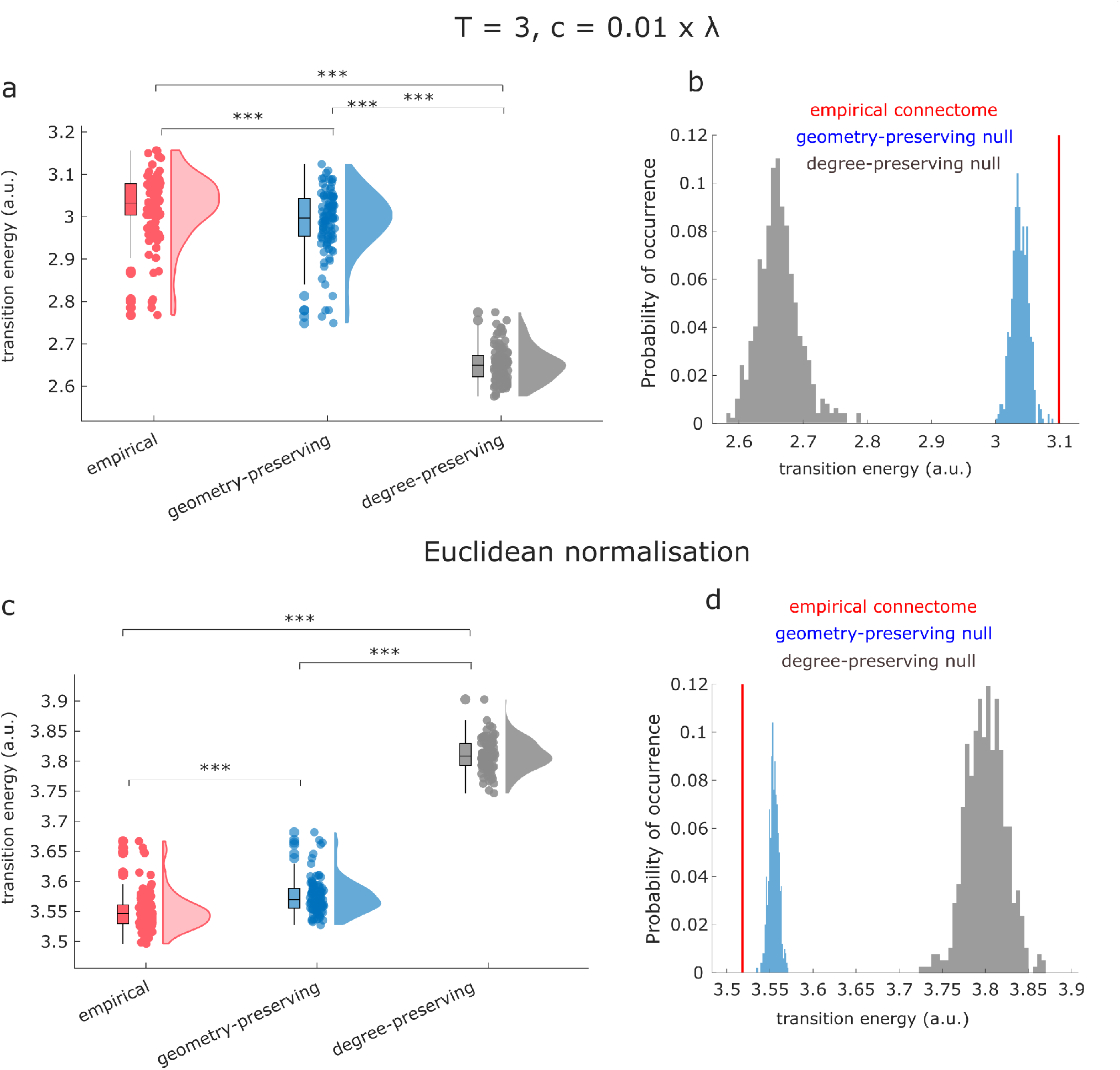
Role of network topology in supporting transitions, for alternative operationalisations of network control theory. **(a,b)** Subject-wise (a) and group-level (b) average transition energy for the empirical human connectome (red) and for degree-preserving (grey) and degree- and cost-preserving null models (blue), for network control with time horizon *T* = 3 and network normalisation factor *c* = 0.01 × |λ(*A*)_max_|. **(c,d)** Subject-wise (c) and group-level (d) average transition energy for the empirical human connectome (red) and for degree-preserving (grey) and degree- and cost-preserving null models (blue), for network control with all NeuroSynth maps normalised to unit Euclidean norm.

**Figure S7.**
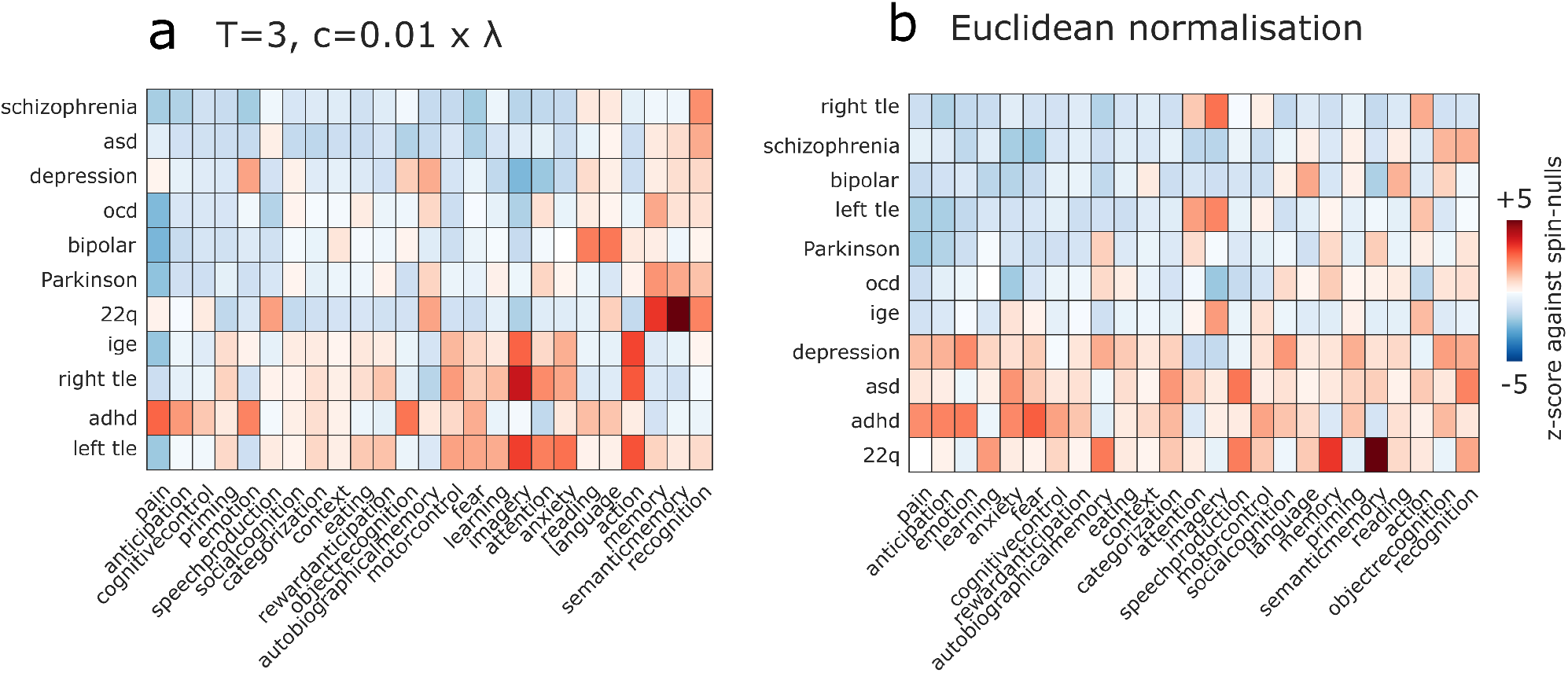
Effects of disease-associated cortical atrophy, for alternative operationalisations of network control theory. Heatmaps show how each disease reshapes the average transition energy required to reach a given cognitive topography from all other cognitive topographies, as a *z*-score against a null distribution of randomly rotated maps with preserved spatial autocorrelation and the same increases and decreases in cortical thickness, but occurring at different neuroanatomical locations. **(a)** For network control with time horizon *T* = 3 and network normalisation factor *c* = 0.01 × |λ(*A*)_max_|; **(b)** for network control with all NeuroSynth maps normalised to unit Euclidean norm. adhd = attention deficit/hyperactivity disorder; asd = autistic spectrum disorder; ocd = obsessive-compulsive disorder; ige = idiopathic generalised epilepsy; right tle = right temporal lobe epilepsy; left tle = left temporal lobe epilepsy.

**Figure S8.**
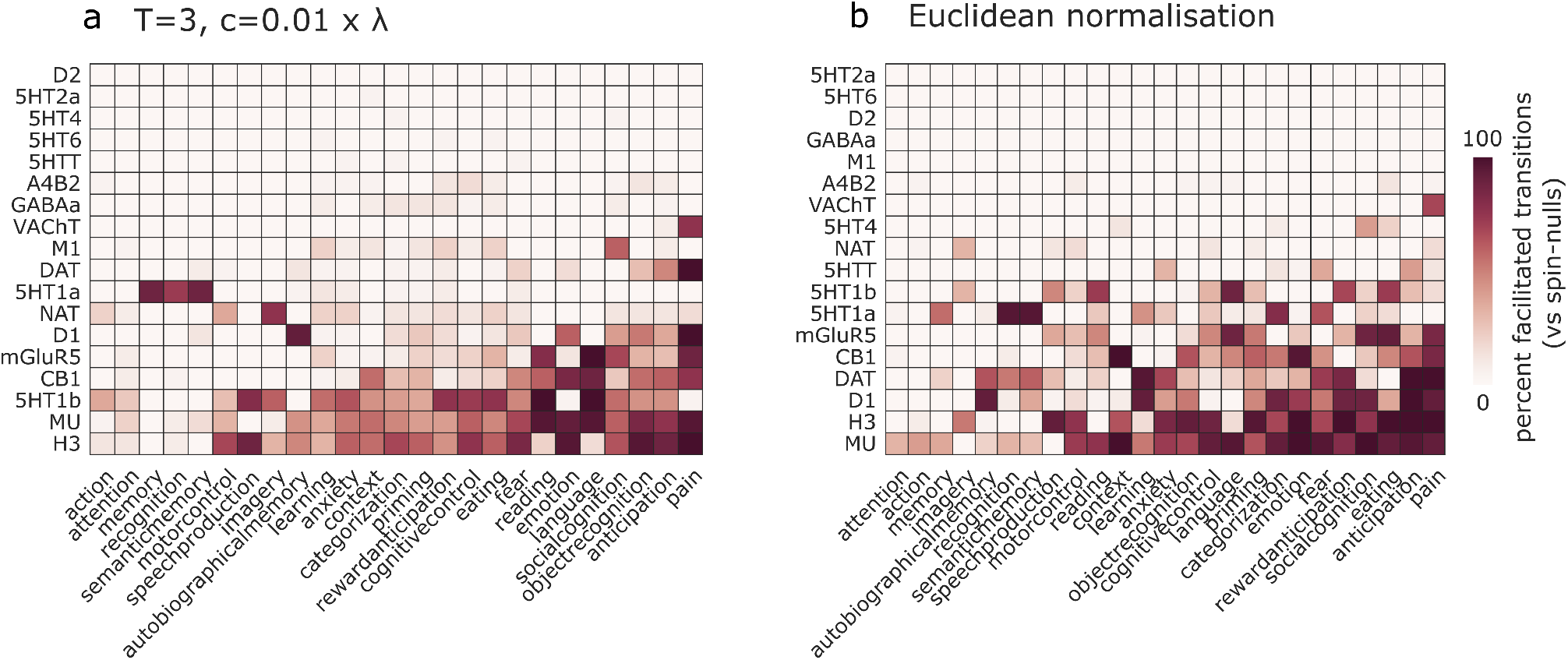
How neurotransmitter systems can reshape the energy landscape of the human brain, for alternative operationalisations of network control theory. Heatmaps show how each receptor/transporter reshapes the average cost of reaching a given cognitive brain state from all other states, as a percentage of transitions to each state that are facilitated, when compared against a null distribution of randomly rotated maps with preserved spatial autocorrelation and the same receptor/transporter density levels, but occurring at different neuroanatomical locations. **(a)** For network control with time horizon time horizon *T* = 3 and network normalisation factor *c* = 0.01 × |λ(*A*)_max_|; **(b)** for network control with all NeuroSynth maps normalised to unit Euclidean norm.

**Figure S9.**
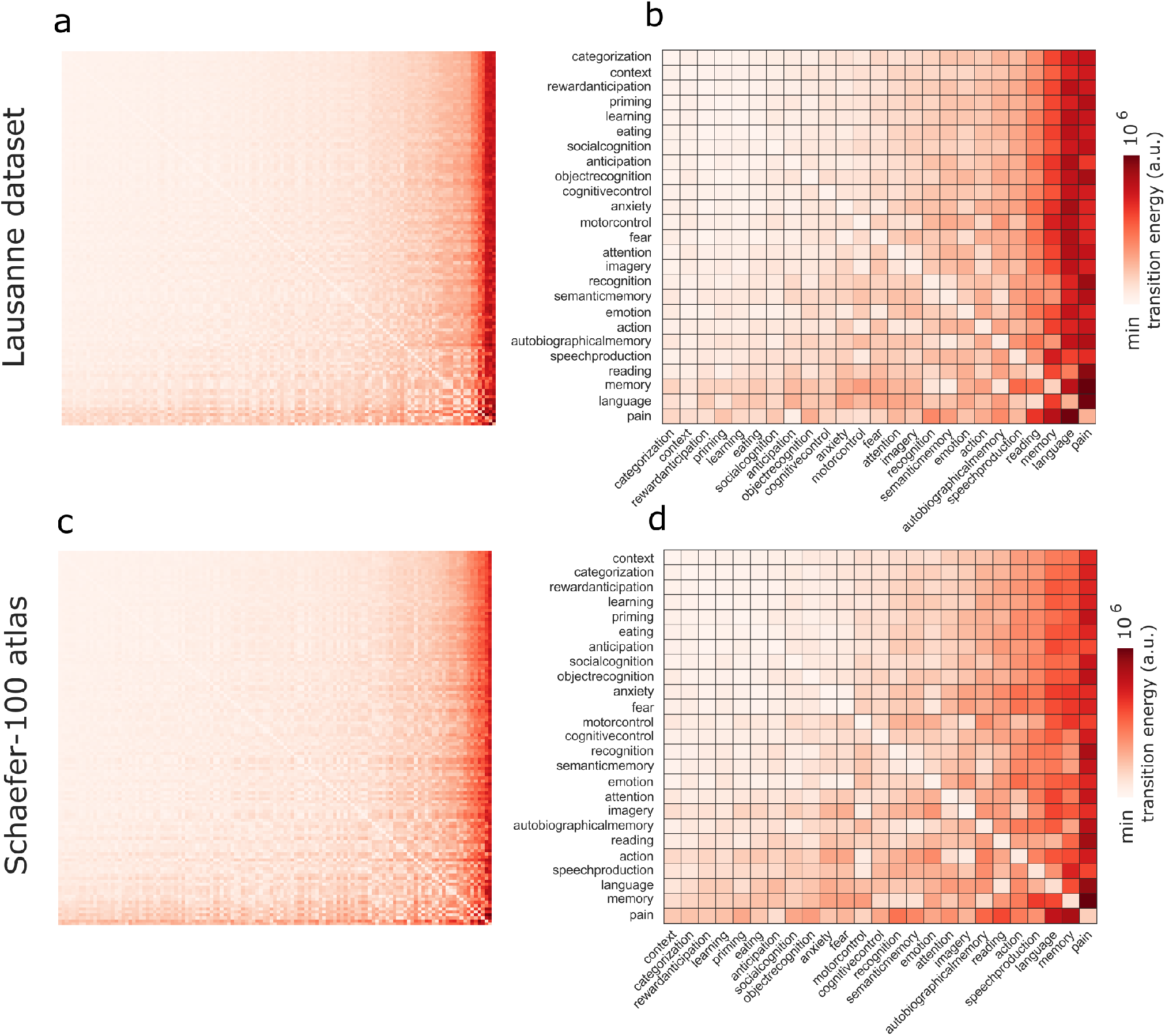
Transition energies for alternative connectome dataset and alternative cortical parcellation. **(a,b)** Transition energy between each pair of 123 cognitive topographies from NeuroSynth (a) and the reduced set of 25 NeuroSynth terms (b), for the Lausanne DSI dataset. Rows indicate source states, columns indicate target states. **(c,d)** Transition energy between each pair of 123 cognitive topographies from NeuroSynth (c) and the reduced set of 25 NeuroSynth terms (d), for Human Connectome Project data parcellated using the Schaefer-100 cortical atlas.

**Figure S10.**
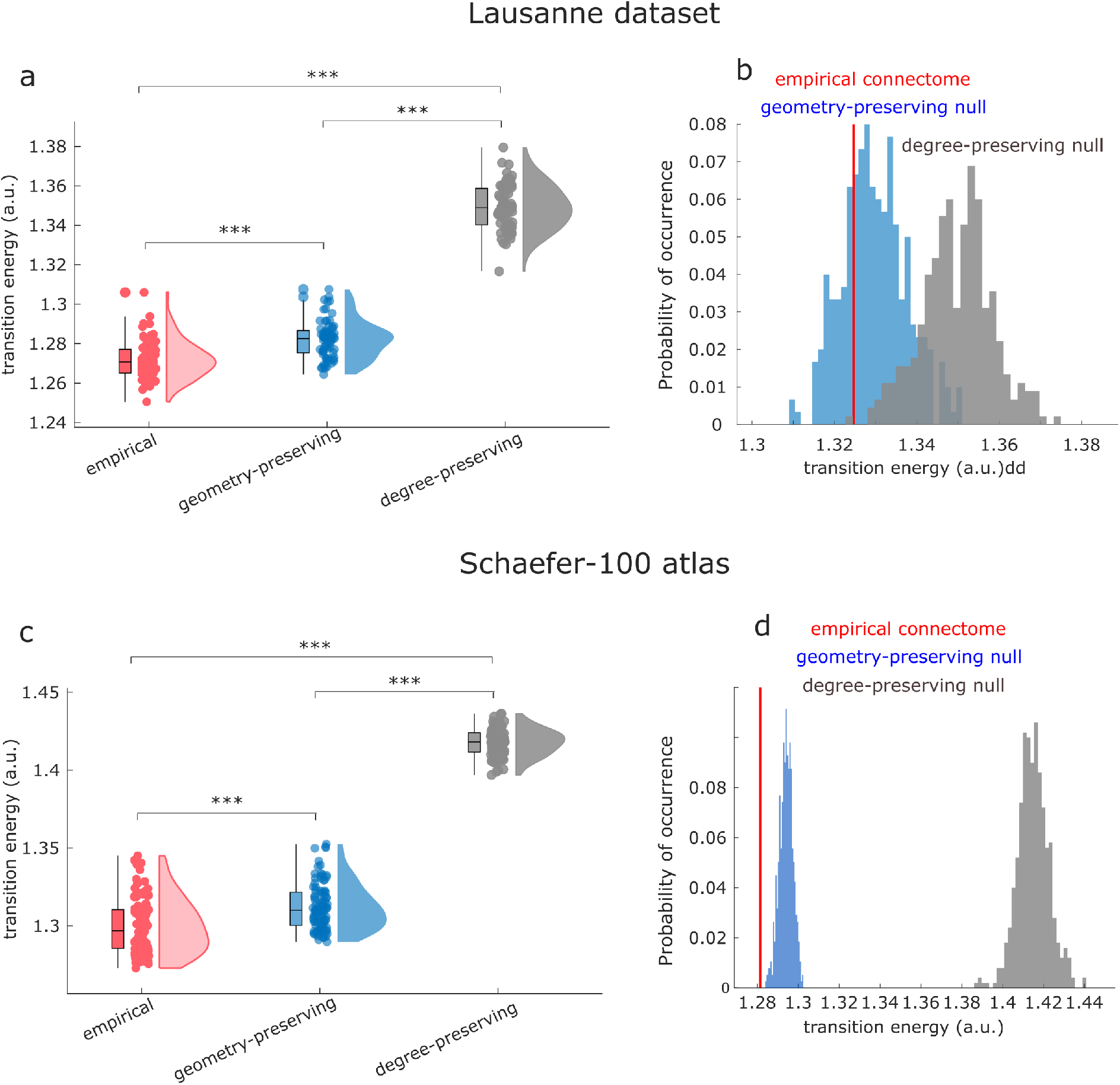
Role of network topology in supporting transitions, for alternative connectome dataset and alternative cortical parcellation. **(a,b)** Subject-wise (a) and group-level (b) average transition energy for the empirical human connectome (red) and for degree-preserving (grey) and degree- and cost-preserving null models (blue), for the Lausanne DSI dataset. **(c,d)** Subject-wise (c) and group-level (d) average transition energy for the empirical human connectome (red) and for degree-preserving (grey) and degree- and cost-preserving null models (blue), for Human Connectome Project data parcellated using the Schaefer-100 cortical atlas.

**Figure S11.**
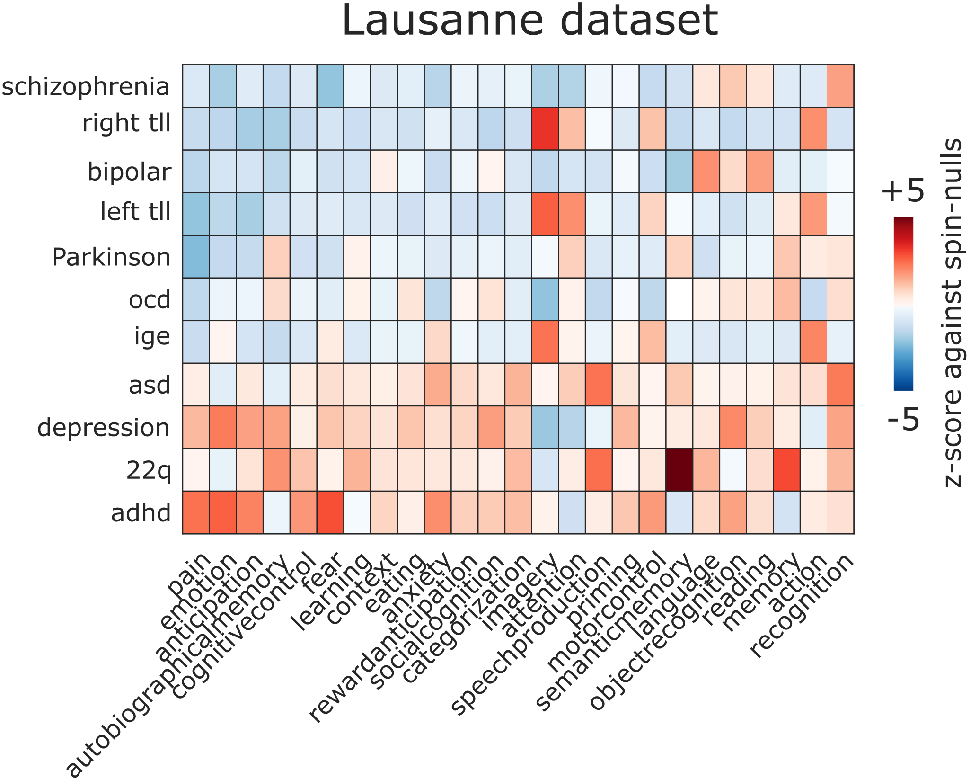
Effects of disease-associated cortical atrophy, for alternative connectome dataset and alternative cortical parcellation. Heatmaps show how each disease reshapes the average transition energy required to reach a given cognitive topography from all other cognitive topographies, as a *z*-score against a null distribution of randomly rotated maps with preserved spatial autocorrelation and the same increases and decreases in cortical thickness, but occurring at different neuroanatomical locations, for the Lausanne DSI dataset. adhd = attention deficit/hyperactivity disorder; asd = autistic spectrum disorder; ocd = obsessivecompulsive disorder; ige = idiopathic generalised epilepsy; right tle = right temporal lobe epilepsy; left tle = left temporal lobe epilepsy.

**Figure S12.**
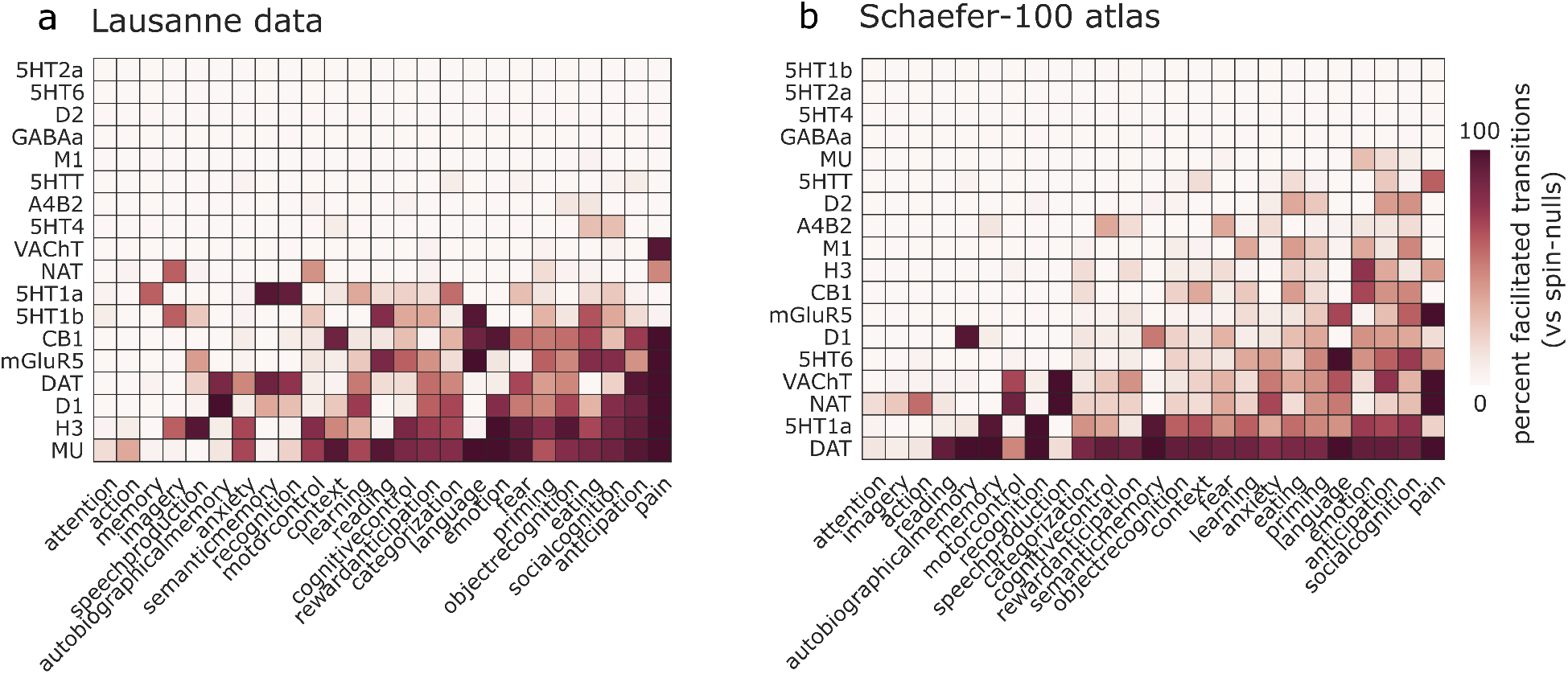
How neurotransmitter systems can reshape the energy landscape of the human brain, for alternative connectome dataset and alternative cortical parcellation. Heatmaps show how each receptor/transporter reshapes the average cost of reaching a given cognitive brain state from all other states, as a percentage of transitions to each state that are facilitated, when compared against a null distribution of randomly rotated maps with preserved spatial autocorrelation and the same receptor/transporter density levels, but occurring at different neuroanatomical locations. **(a)** For the Lausanne DSI dataset; **(b)** for Human Connectome Project data parcellated using the Schaefer-100 cortical atlas.

**Figure S13.**
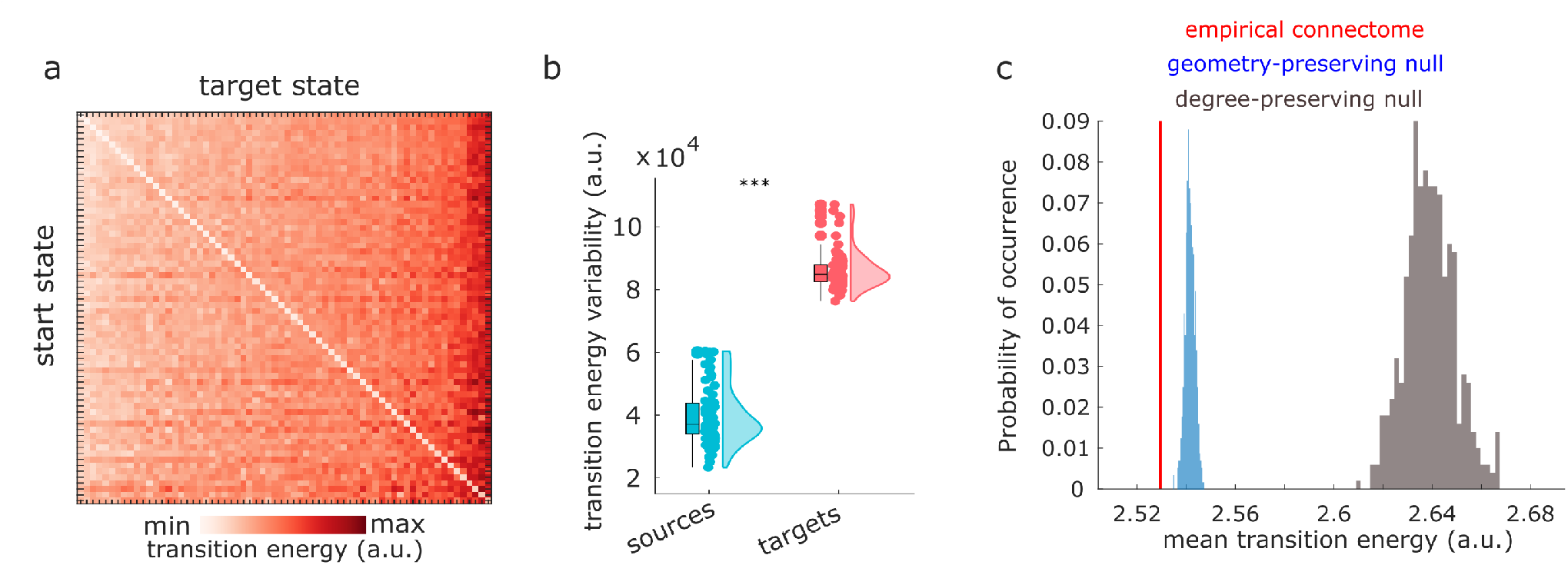
Replication with cognitive topographies defined by BrainMap. **a** Transition energy between each pair of 66 cognitive topographies from BrainMap. **b** Variability (standard deviation) of transition energy is greater along target states than along start states. **c** Degree-preserving randomised null models (grey) and null models that preserve the exact degree sequence and the approximate length distribution (blue) are significantly less favourable than the empirical human connectome (red) to support transitions between cognitive topographies defined by BrainMap.

**TABLE S1.** Subject-level comparison against null networks. Statistical comparison between subject-level overall transition energy distributions, for the empirical human connectome (*N* = 100 Human Connectome Project subjects) and corresponding degree-preserving and degree- and cost-preserving rewired nulls.

|  | Mean1 | SD1 | Mean2 | SD2 | df | t-score | p-value | Effect Size |
| --- | --- | --- | --- | --- | --- | --- | --- | --- |
| empirical vs geometry-preserving | 1.25e+06 | 13400 | 1.26e+06 | 12200 | 99 | -21.49 | <0.001 | -0.77 |
| empirical vs degree-preserving | 1.25e+06 | 13400 | 1.35e+06 | 9160 | 99 | -60.18 | <0.001 | -9.05 |
| degree- vs geometry-preserving | 1.35e+06 | 9160 | 1.26e+06 | 12200 | 99 | 58.92 | <0.001 | 8.7 |

**TABLE S2.** Subject-level results for the Lausanne dataset. Statistical comparison between subject-level overall transition energy distributions, for the empirical human connectome (*N* = 70 subjects from the Lausanne dataset) and corresponding degree-preserving and degree- and cost-preserving rewired nulls.

|  | Mean1 | SD1 | Mean2 | SD2 | df | t-score | p-value | Effect Size |
| --- | --- | --- | --- | --- | --- | --- | --- | --- |
| empirical vs geometry-preserving | 1.27e+06 | 9560 | 1.28e+06 | 9630 | 69 | -15.03 | <0.001 | -1.1 |
| empirical vs degree-preserving | 1.27e+06 | 9560 | 1.35e+06 | 11300 | 69 | -44.91 | <0.001 | -7.3 |
| degree- vs geometry-preserving | 1.35e+06 | 11300 | 1.28e+06 | 9630 | 69 | 36.34 | <0.001 | 6.3 |

**TABLE S3.**
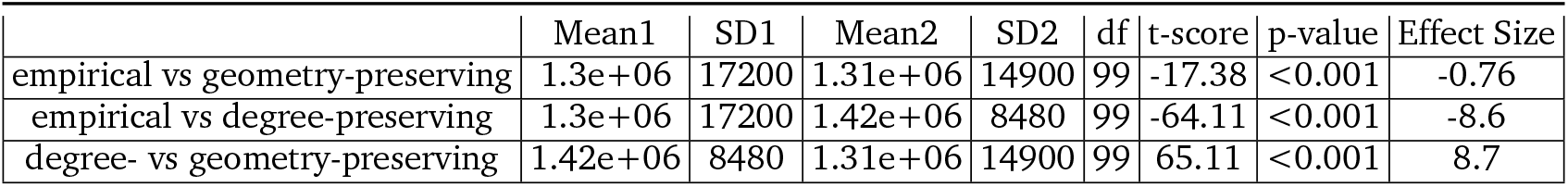
Subject-level results for Schaefer-parcellated data. Statistical comparison between subject-level overall transition energy distributions, for the empirical human connectome (*N* = 100 subjects from the Human Connectome Project) and corresponding degree-preserving and degree- and cost-preserving rewired nulls.

|  | Mean1 | SD1 | Mean2 | SD2 | df | t-score | p-value | Effect Size |
| --- | --- | --- | --- | --- | --- | --- | --- | --- |
| empirical vs geometry-preserving | 1.3e+06 | 17200 | 1.31e+06 | 14900 | 99 | -17.38 | <0.001 | -0.76 |
| empirical vs degree-preserving | 1.3e+06 | 17200 | 1.42e+06 | 8480 | 99 | -64.11 | <0.001 | -8.6 |
| degree- vs geometry-preserving | 1.42e+06 | 8480 | 1.31e+06 | 14900 | 99 | 65.11 | <0.001 | 8.7 |

**TABLE S4.**
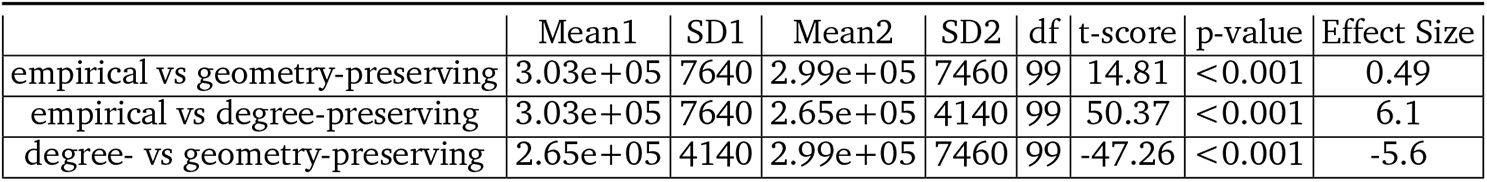
Subject-level results for alternative implementation of network control theory. Statistical comparison between subject-level overall transition energy distributions, for the empirical human connectome (*N* = 100 HCP subjects) and corresponding degree-preserving and degree- and cost-preserving rewired nulls, for network control with time horizon *T* = 3 and network normalisation factor *c* = 0.01 × |λ(*A*)_max_|.

**TABLE S5.**
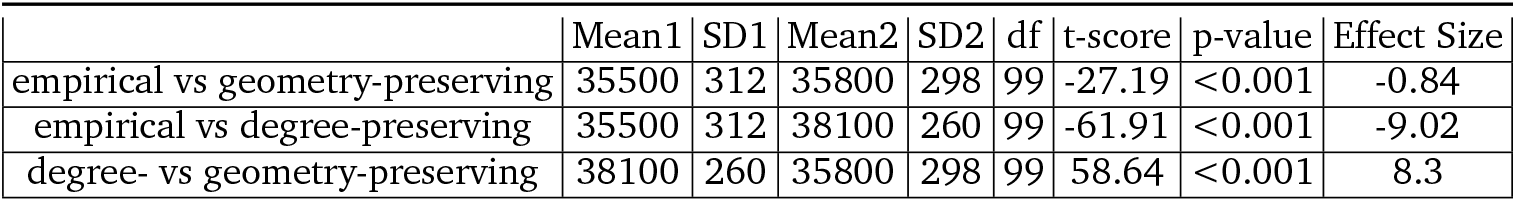
Subject-level results for alternative implementation of network control theory. Statistical comparison between subject-level overall transition energy distributions, for the empirical human connectome (*N* = 100 HCP subjects) and corresponding degree-preserving and degree- and geometry-preserving rewired nulls, for network control with NeuroSynth maps normalised to unit Euclidean norm

